## Supplementary Information for "Iron rescues glucose-mediated photosynthesis repression during lipid accumulation in the green alga *Chromochloris zofingiensis*"

Table of Contents

**Evolutionary comparisons of highest-in-heterotrophy proteins** 1.
**Detailed Supplementary Methods** 2.
***Proteomic Mass-Spectrometry*** 2.
*Protein extractions and processing for proteomics* 2.
*TMT isobaric tag labeling* 3.
*High pH RP C-18 fractionation* 3.
***Lipid extraction and thin layer chromatography.*** 3.
***Carotenoid and chlorophyll detection by high performance liquid chromatography.*** 4.
**Supplementary Figures (1-11)** 5.
**Supplementary Table 1** 14.
**Supplementary References** 15.

**Evolutionary comparisons of highest-in-heterotrophy proteins**

Loss of photosynthesis was coupled with loss of several known members of the photosynthetic proteins. However, besides loss of photosynthesis, we lacked an understanding of the predominant biochemistries and physiologies that were upregulated during green algal heterotrophy. Therefore, we used bioinformatic functional predictions and cross-species comparisons of the highest-in-heterotrophy proteins to show unique physiology and iron priorities in *C. zofingiensis.* For example, cobalamin-independent methionine synthase (METE) is stringently highest-in-heterotrophy. However, its upregulation is unlikely to be proteinogenic as several other amino acid biosynthesis family proteins are generally low or lowest-in-heterotrophy (e.g. glutamate synthases (Cz08g04180, Cz10g25220), a glutamine synthetase (Cz12g28120), alanine aminotransferases (Cz11g04020, Cz11g2410)). In addition, GO analysis of highest-in-heterotrophy proteins showed enrichment for “thiamine biosynthetic process” (GO:0009228, $p$ = 0.0014). The enzymes represented by these GO categories include THI1/4 proteins, which react with glycine, NAD+, and a sulfur donor to make the thiazole moiety of thiamine**,** and THIC, which synthesizes the pyrimidine moiety.

*C. zofingiensis* unexpectedly contains three genes encoding THI1/4 whereas other photosynthetic organisms usually have one copy (Supplementary Fig. 10a). All three THI1/4 proteins and THIC were distinguished by unique peptides and were highly abundant in heterotrophy (Supplementary Fig. 10b). Both THIC and THI1/4 contain iron cofactors (4Fe-4S cluster and Fe cation, respectively), which would normally suggest their depletion in −Fe. In fact, we found THIC and THI1/4 were depleted in −Fe in two *C. reinhardtii* iron-deficiency proteomes[1,2] (Supplementary Fig. 10c,d). METE, which contains a zinc cofactor, was also depleted by −Fe in *C. reinhardtii*[1,2]. This suggests METE and thiamine biosynthesis enzyme upregulation in *C. zofingiensis* is a distinct heterotrophy feature in contrast to *C. reinhardtii* Fe deficiency. In addition, both gene duplication and upregulation in Fe-dependent heterotrophy of all iron cofactor-containing THI players suggests *C. zofingiensis* displays an evolutionary novel iron sink priority in WT−Fe+Glc (see Fig. 9).

The example of unique highest-in-heterotrophy proteins show how iron priorities during deficiency may diverge according to an organism's environment and evolutionary programming. The iron-cofactor proteins for thiazole biosynthesis and PAO5 were uniquely upregulated in WT−Fe+Glc and could be iron sinks when several other iron-rich proteins are depleted in heterotrophy. In particular, the THI1/4 family has duplicated to three copies in *C. zofingiensis,* all of which are induced in heterotrophy (Fig. 9). This protein family is known to be metabolically costly due to catalyzing only a single reaction turnover per protein[3]. It is surprising that *C. zofingiensis* upregulates these biosynthetic proteins in −Fe+Glc despite their metabolic and iron costs, especially if the THI1/4 genomic duplications evolutionarily increased this protein's concentration or led to neo-functionalization. Further research may unveil the importance of the thiazole moiety to heterotrophic metabolism, which could include novel enzymatic uses for thiazole-derived products in *C. zofingiensis*.

#### Detailed Supplementary Methods

##### ***Proteomic Mass-Spectrometry***

*Protein extractions and processing for proteomics*

For protein extraction, each cell pellet was resuspended in 200 µl of $H_2O$ and transferred to 2 mL pre-filled Micro-Organism Lysing Mix glass bead tubes and disrupted in a Bead Ruptor Elite bead mill homogenizer (OMNI International, Kennesaw, GA) at speed 5.5 m/s for 45 s. After bead beating, the lysate was immediately placed on ice and then centrifuged at 1,000 *x g* for 10 mins at 4°C. To separate the proteins, metabolites and lipids, 1 mL cold (-20°C) 2:1 chloroform:methanol (v/v) was pipetted into a chloroform-compatible 2 mL Sorenson MulTI™ SafeSeal™ microcentrifuge tube (Sorenson Bioscience, Salt Lake City, UT) on ice. The 200 µl of sample homogenate was then added to the Sorenson tube and vigorously vortexed. The sample was placed on ice for 5 min and then vortexed for 10 s followed by centrifugation at 10,000 *x g* for 10 min at 4°C. The protein interface had 1 mL of cold 100% methanol added to each sample, was vortexed and centrifuged again at 10,000 *x g* for 10 min at 4°C to pellet the protein. The methanol was then decanted off, and the samples were placed open in a fume hood to dry for ~10 min.

The protein pellet was dissolved in 200 µl of 8 M urea and vortexed into solution. A bicinchoninic acid (BCA) assay (Thermo Scientific, Waltham, MA USA) was performed to determine protein concentration. Following the assay, 10 mM dithiothreitol (DTT) was added, and the samples were incubated at 60°C for 30 min with constant shaking at 800 rpm. Samples were then diluted 8-fold in preparation for trypsin digestion. 100 mM $NH_4HCO_3$, 1 mM $CaCl_2$ and sequencing-grade modified porcine trypsin (Promega, Madison, WI) were added to all protein samples at a 1:50 (w/w) trypsin-to-protein ratio for 3 h at 37˚C with constant shaking at 450 rpm. Digested samples were desalted using a 4-probe positive pressure Gilson GX-274 ASPEC™ system (Gilson Inc., Middleton, WI) with Discovery C18 100 mg/1 mL solid phase extraction tubes (Supelco, St. Louis, MO), using the following protocol: 3 mL of methanol was added for conditioning followed by 2 mL of 0.1% trifluoroacetic acid (TFA) in $H_2O$. The samples were then loaded onto each column followed by 4 mL of 95:5: $H_2O$: acetonitrile, 0.1% TFA. Samples were eluted with 1 mL 80:20 acetonitrile:$H_2O$, 0.1% TFA. The samples were concentrated down to ~100µL using a SpeedVac, and a final BCA assay was performed to determine the peptide concentration. An equal mass of each sample was aliquoted into fresh centrifuge tubes, dried completely in a SpeedVac and then stored at -80°C until isobaric labeling.

*TMT isobaric tag labeling*

Each sample was diluted in 500 mM HEPES, pH 8.5 to a concentration of 5 µg/µl and labeled using amine-reactive Thermo Scientific Tandem Mass Tag (TMT10) Isobaric Mass Tagging Kits (Thermo Scientific, Rockford, IL) according to the manufacturer's instructions. Briefly, 250 µL of anhydrous acetonitrile was added to each 5 mg reagent, vortexed and allowed to dissolve for 5 min with occasional vortexing. Reagents were then added to the samples and incubated for 1 h at RT with shaking at 400 rpm. Each sample was diluted to 2.5 µg/µl with 20% acetonitrile, and the reaction was quenched by adding 8 µL of 5% hydroxylamine to the sample with incubation for 15 min at RT with shaking at 400 rpm. Samples within each set were combined and completely dried in the SpeedVac. Each sample was cleaned using C18 50 mg/1 mL solid phase extraction tubes as described above and again assayed with BCA to determine the final peptide concentration.

*High pH RP C-18 fractionation*

TMT samples were diluted to a volume of 900 µL with 10 mM ammonium formate buffer (pH 10.0), and resolved on a XBridge C18, 250x4.6 mm, 5 µm with 4.6x20 mm guard column (Waters, Milford, MA). Separations were performed at 0.5 mL/min using an Agilent 1100 series HPLC system (Agilent Technologies, Santa Clara, CA) with mobile phases (A) 10 mM ammonium formate, pH 10.0 and (B) 10 mM ammonium formate, pH 10.0/acetonitrile (10:90). The gradient was adjusted from 100% A to 95% A over the first 10 min, 95% A to 65% A over minutes 10 to 70, 65% A to 30% A over minutes 70 to 85, maintained at 30% A over minutes 85 to 95, re-equilibrated with 100% A over minutes 95 to 105, and held at 100% A until minute 120. Fractions were collected every 1.25 minutes (96 fractions over the entire gradient). Every other row was concatenated into 24 fractions and dried completely, and 25 µL of 25 mM ammonium bicarbonate was added to each fraction for storage at -20°C until LC-MS/MS analysis.

***Lipid extraction and thin layer chromatography.***

The procedures for visualizing lipids by TLC were based off previous methods[4]. Cell pellets from 2mL of culture were flash frozen. Cells were lysed in 2 mL screw-cap tubes with Lysing Matrix D beads through a MP Fastprep-24™ 5G bead beater (settings 6.5 m/s, 60s, 2 rounds) with dry ice in the CoolPrep™ adapter. The frozen cell pellets were given 0.5 mL (except the high biomass WT+Fe+Glc sample, where extraction volumes were doubled) of 2:1 chloroform:methanol solution with 0.01% butylated hydroxytoluene and were vortexed for 5 min. 133 µL (266 for WT+Fe+Glc) of 0.73% (w/v) NaCl was added to the cultures, which were centrifuged for 2min at 14,000 rpm (20317 RCF). The lower solvent with extracted lipids was transferred Eppendorf tubes. The volume of extracted lipids added to Supelco TLC Silica gel 60 $F_{254}$ (1.05715.0001) pretreated with developing solution was adjusted so that the same culture biomass (2.51 x $10^7$ µ$m^3$) per extraction solution was loaded on plate. 2.5 µL of lipid extracted olive oil (25 µL Good & Gather, extracted in 0.5 mL of 2:1 chloroform:methanol solution as above) was also loaded as a TAG standard. The plate was developed with hexane:diethyl ether:acetic acid (91:30:1.3) which shows strong separation of TAGs [8].The plate was lightly sprayed with 25% $H_2SO_4$ in 50% Ethanol with a Aldich[R] flask-type sprayer (75 mL), dried for 10 minutes, than charred in a ~100°C oven until the lipid bands were prominent. The TLC plate was photographed while being illuminated by Fotodyne UV Transilluminator.

***Carotenoid and chlorophyll detection by high performance liquid chromatography***

The protocol was conducted as had been done previously in *C. zofingiensis*[4-6]. Briefly, 5 mL of culture at the 84 h sampling time point had growth measurements were spun down, the supernatant was removed, and remaining cell pellet was flash frozen. Samples were resuspended in 100% HPLC-grade acetone and homogenized with Matrix D beads in a FastPrep-24 5G™High-Speed Homogenizer (6.5 m s–1 for 2 x 60 s, MP Biomedical). The pigmented supernatant was collected after centrifugation (2 min, 1500 g, 4°C) and the sample was repeated until the cell pellet was white and supernatant was mostly clear towards the last extractions. All steps were held on ice, in the dark as much as possible. Sample runs through the HPLC were conducted by a previously described chlorophyll and carotenoid quantification protocol[4-7] on an Agilent 1100 HPLC system.

### Supplementary Figures

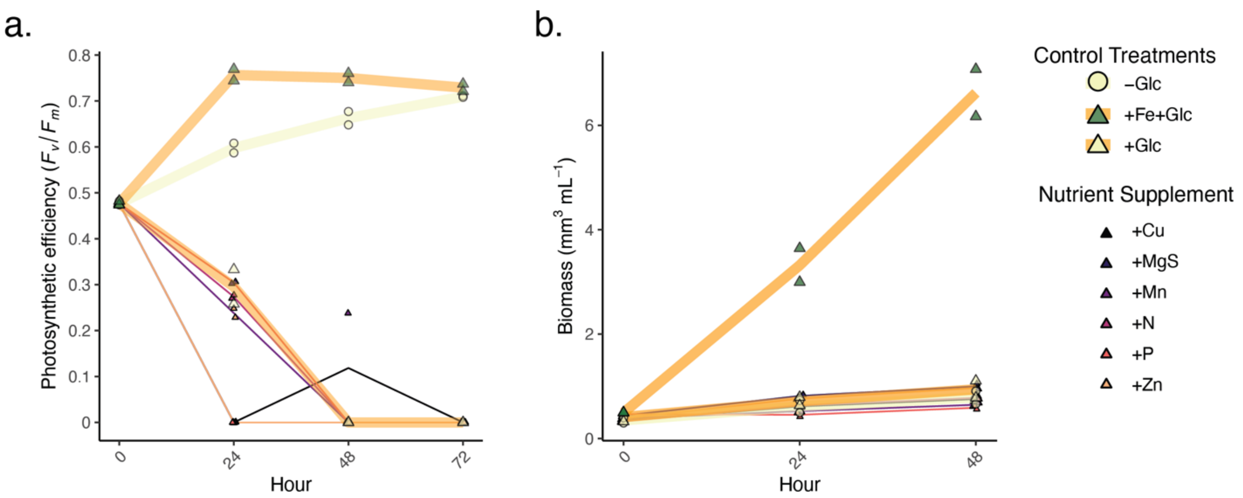


**Supplementary Fig. 1 Photosynthesis rescue and biomass gain in +Glc only occurs from +Fe.**
The nutrient impacts of adding copper (2 µM $Cu^{2+}$ EDTA), manganese (6 µM Mn-EDTA), (2.5 µM $Zn^{2+}$- EDTA) (Concentrations from Kropat et al. 2011), nitrogen (10 mM $NaNO_3$), magnesium and sulfur (1 mM $MgSO_4$), and phosphorus (2 mM $K_2PO_4/KH_2PO_4$) alongside glucose were compared to the control treatments of −Glc, +Glc, and +Fe+Glc (thick lines, larger datapoints). **a.** Maximum quantum efficiency of PSII *($F_v/F_m$)* with nutrient supplements over 72 h. **b.** Biomass (per mL) with nutrient supplements over 72 h. *n* = 2 for each treatment.

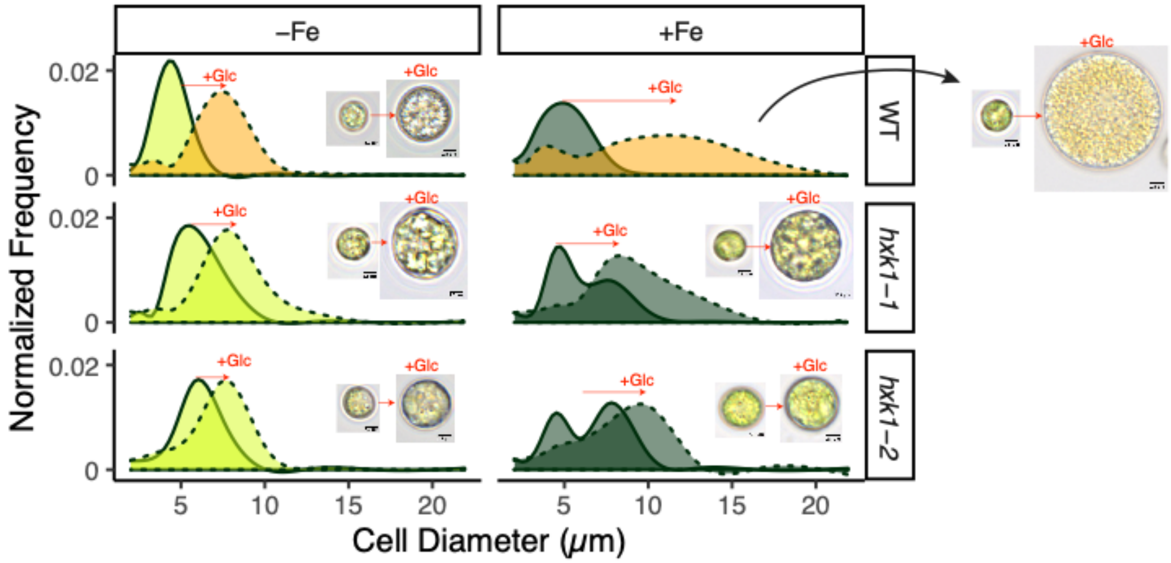


**Supplementary Fig. 2 Glucose increases cell size of all strains and Fe conditions.**
Distribution of cell sizes are normalized by their cell density for comparison. The dotted line (+Glc) versus the solid line (−Glc) shows how all strains respond to glucose to become large cells. The mixotrophic WT+Fe+Glc cells have the largest mean cell size but also the largest range in cell size. Corresponding light microscopy images have the same size scalebar (2.5 µm), and mostly close to median cells for each of the 12 conditions. However, one of the larger cells was chosen for WT+Fe+Glc to show maximum size range. Cell clumping did not occur in any culture and does not contribute to cell size distributions.

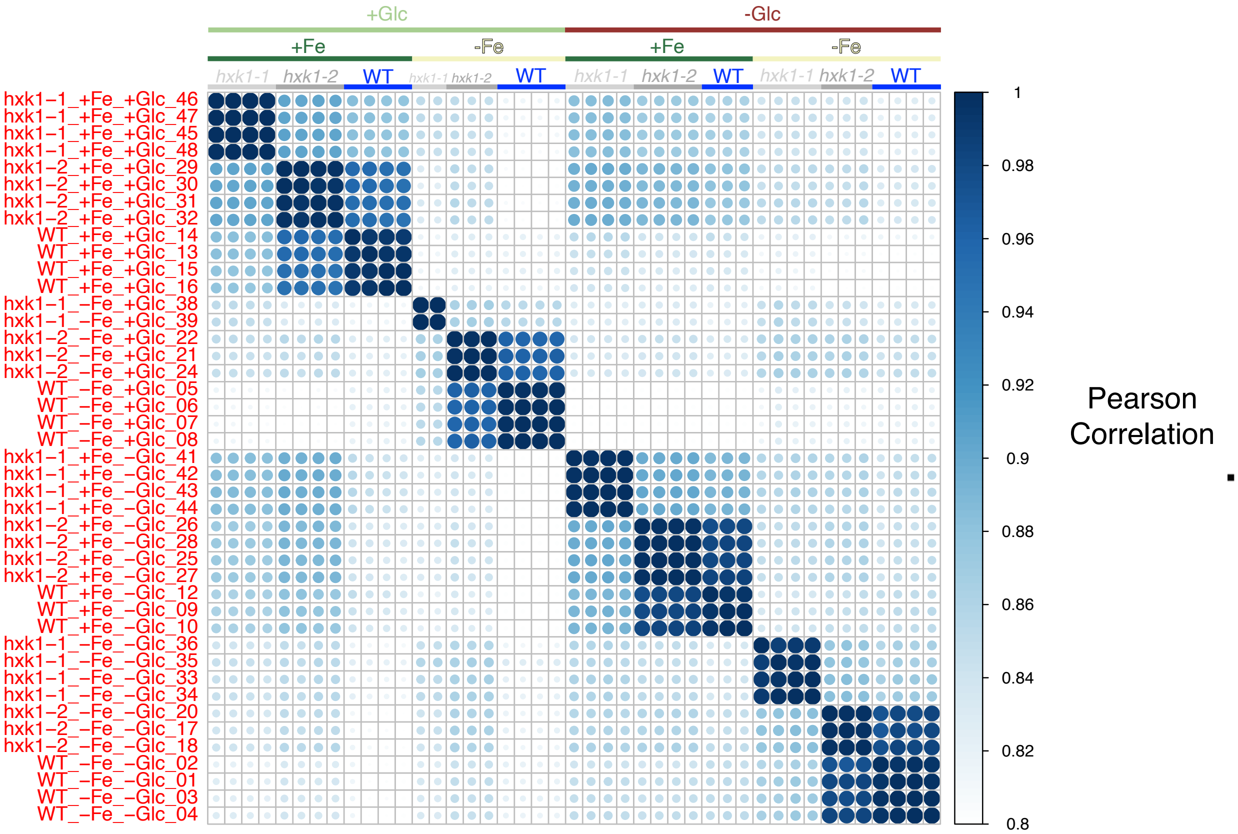


**Supplementary Fig. 3 High condition global proteome replicability in autocorrelation plot.**
Correlation plot (R's corrplot package) of each sample. Each row and column represent a condition and the blue scale and circle size represent the Pearson correlation each protein (detected in all conditions) of two samples. A color key on top indicates condition variables (Yellow: −Fe, Dark Green: +Fe; Light Green: −Glc, Brown: +Glc; Grey: *hxk1-2*, Blue:WT). Individual sample IDs are labelled in *y*-axis. The Pearson correlation scale was narrowed to 0.8-1 to focus on the subtle changes in correlation.

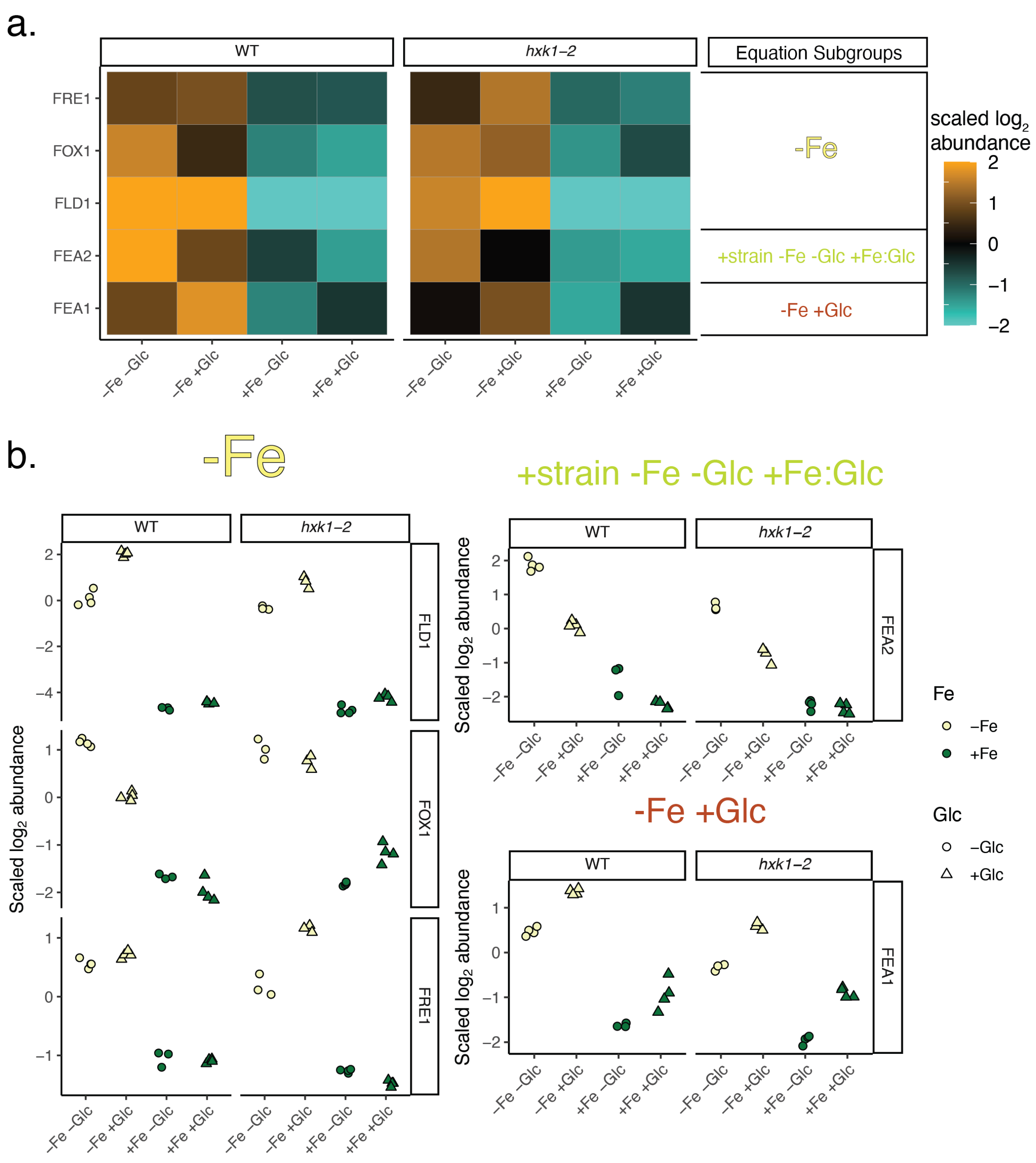


**Supplementary Figure 4. Categorization of Fe deficiency biomarkers shows extent of sensitivity of linear model categorization.**

**a.** Proteomics heatmap of Fe deficiency biomarkers shows general upregulation in −Fe conditions. Equation subgroups as the result of the statistical categorization pipeline for each biomarker belongs to are labelled in the right. **b.** Raw plots of each replicate protein abundance are additionally labelled to show nuances of the dominant linear model assignment. The five Fe deficiency biomarkers are well captured to be upregulated in −Fe conditions, although the linear model filtering pipeline captures Glc and strain impacts as well. Flavodoxin 1 (FLD1, highest upregulated in −Fe), multicopper ferroxidase 1 (FOX1) and ferric reductase 1 (FRE1) all were placed in the −Fe group, where protein distributions are dominated by the increase in −Fe. Fe-assimilating protein 1 (FEA1) is categorized into

“−Fe+Glc” equation subgroup, where its activity increase −Fe deficient but also in response +Glc independent of Fe. FEA2 was categorized into the subgroup “+strain −Fe −Glc +Fe:Glc”. While repressed in +Fe, this protein has higher abundance in WT−Fe−Glc than *hxk1-2*−Fe−Glc or WT−Fe+Glc.

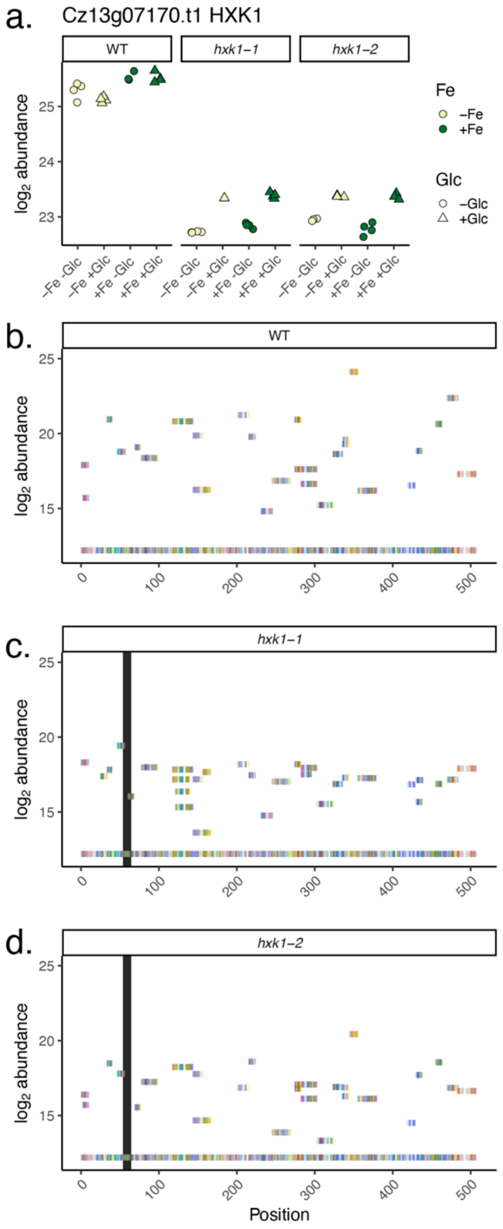


**Supplementary Fig. 5 Low abundance of HXK1 is detected in *hxk1* strains.**
**a.** Relative protein abundance of HXK1 across strains (n=3-4, individual data points shown). Glc and Fe had minimal impact on abundance at this 84 h timepoint. **b-d**. Unique Peptides Aligned to the HXK1 protein sequence. For each strain, each rectangle represents uniquely detected peptides and their aligned to their position on the protein sequence (*x*-axis). *y*-axis position signifies relative abundance of each peptide and each amino acid is colored according to RasMol (2.7.5) color schemes. The black rectangle in the hxk1 strains signifies the site where the C471CG insertion is supposed to confer a frame shift (AA position 55) to the mutant protein’s predicted early stop (AA position 65). Most peptides downstream align to downstream of the frame shift and stop. Plots **b-d** shows the mean unique peptide abundance within +Fe−Glc conditions per strain, but peptides aligned downstream of the frameshift mutation occur in all conditions.

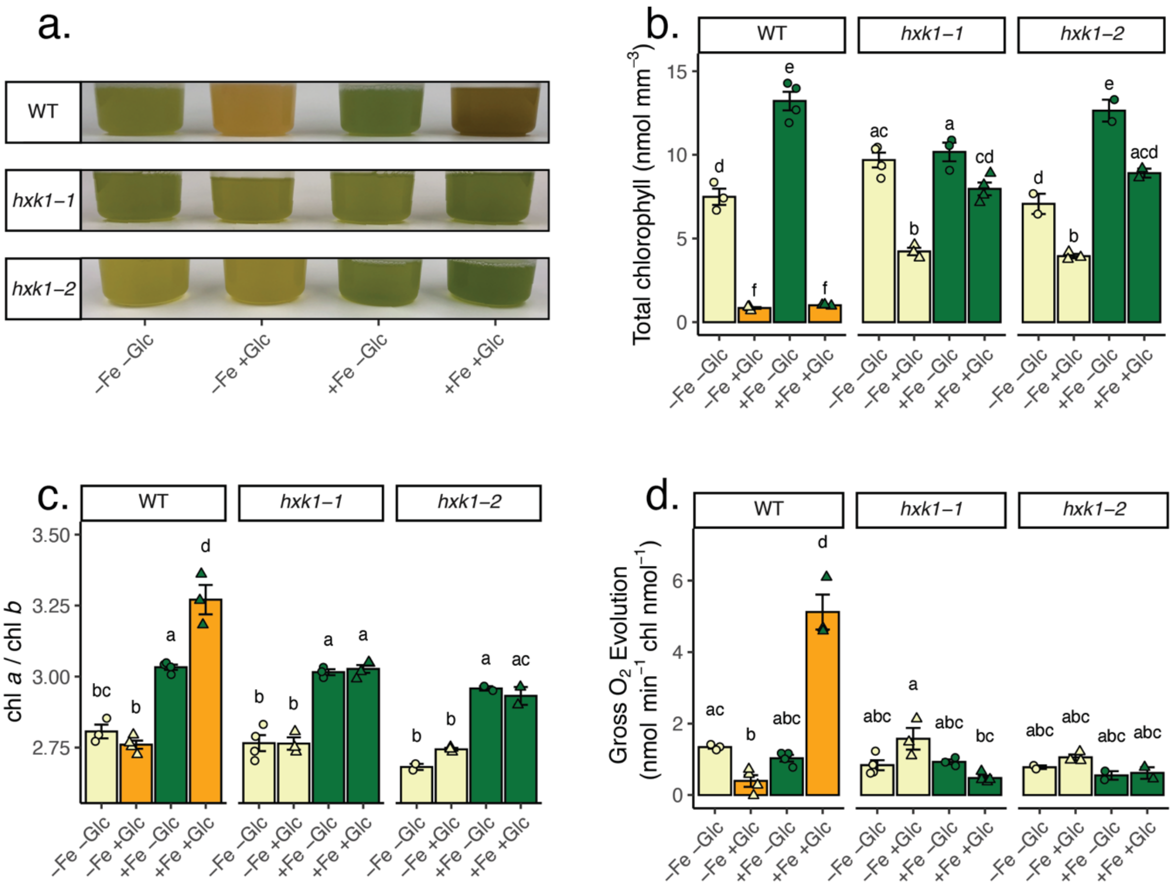


**Supplementary Fig. 6 Mixotrophic (+Fe+Glc) samples maintain reaction center chlorophyll.**
**a.** Photograph of representative cultures at 84 h, right before sampling for HPLC. **b.** Total chlorophyll (*a* + *b*) normalized to $mm^3$ of volumetric biomass. Compare to normalized per cell in Figure 4c. **c.** The chlorophyll a/ chlorophyll *b* ratio of the 12 conditions. Chlorophyll *a* is part of core reaction centers while both chl *a* and chl *b* are part of periphery light harvesting complexes. **d.** Gross oxygen evolution normalized to total chlorophyll. For all graphs, data represent means $\pm$SE, with individual points $n$ = 2-4. Statistical comparisons are applied by Tukey's HSD with adjusted $p$-value = 0.05 and represented by letters above bars.

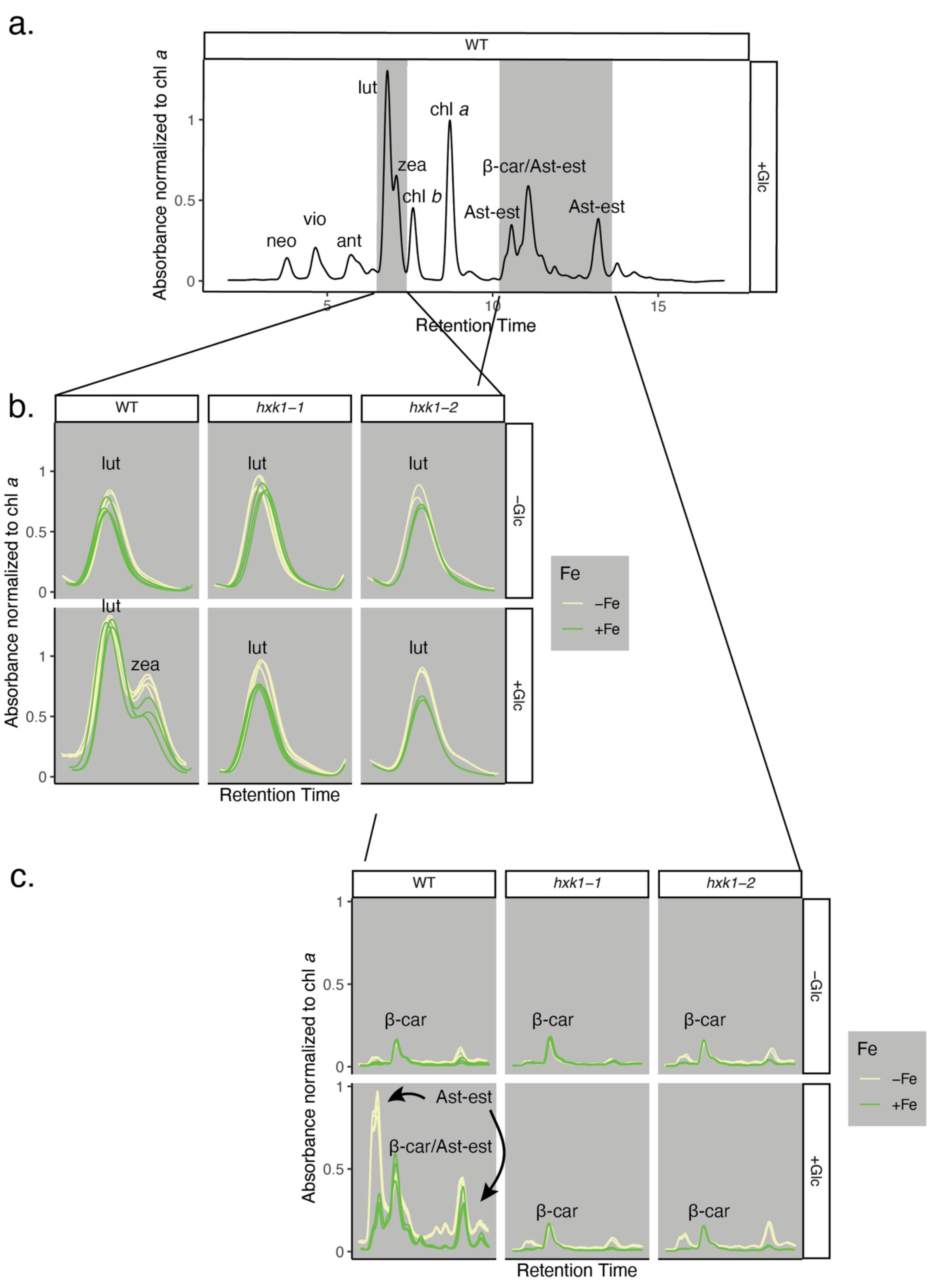


**Supplementary Fig. 7 Zeaxanthin and astaxanthin accumulate in WT+Glc.**

**a.** Representative HPLC chromatograms of chlorophyll and carotenoids normalized to the peak absorbance of chlorophyll *a*. The known pigments in the chromatogram are neoxanthin (neo), violaxanthin (vio), antheraxanthin (ant), lutein (lut), zeaxanthin (zea), chlorophyll *b* (chl *b*), chlorophyll *a* (chl *a*), β-carotene (β-car), and esterified astaxanthin (Ast-est). This representative sample is WT+Fe+Glc with key pigments filled in grey, **b.** lutein and zeaxanthin and **c.** esterified astaxanthin **b.** Grid of lut and zea zoomed in retention times. The zeaxanthin "shoulder" peak only appears in the WT+Glc grid, both in +Fe (green) and −Fe (cream yellow). **c.** Esterified-astaxanthin is mostly found in two peaks or three peaks due to astaxanthin being in mono-esterified or di-esterified forms, which have different retention times[6]. These signals accumulate in WT−Fe+Glc and WT+Fe+Glc greater than any other condition. β-car is found in all conditions and likely overlaps with esterified-astaxanthin in the WT+Glc conditions. Each individual line is biological replicate from 84 h ($n$ = 2-4).

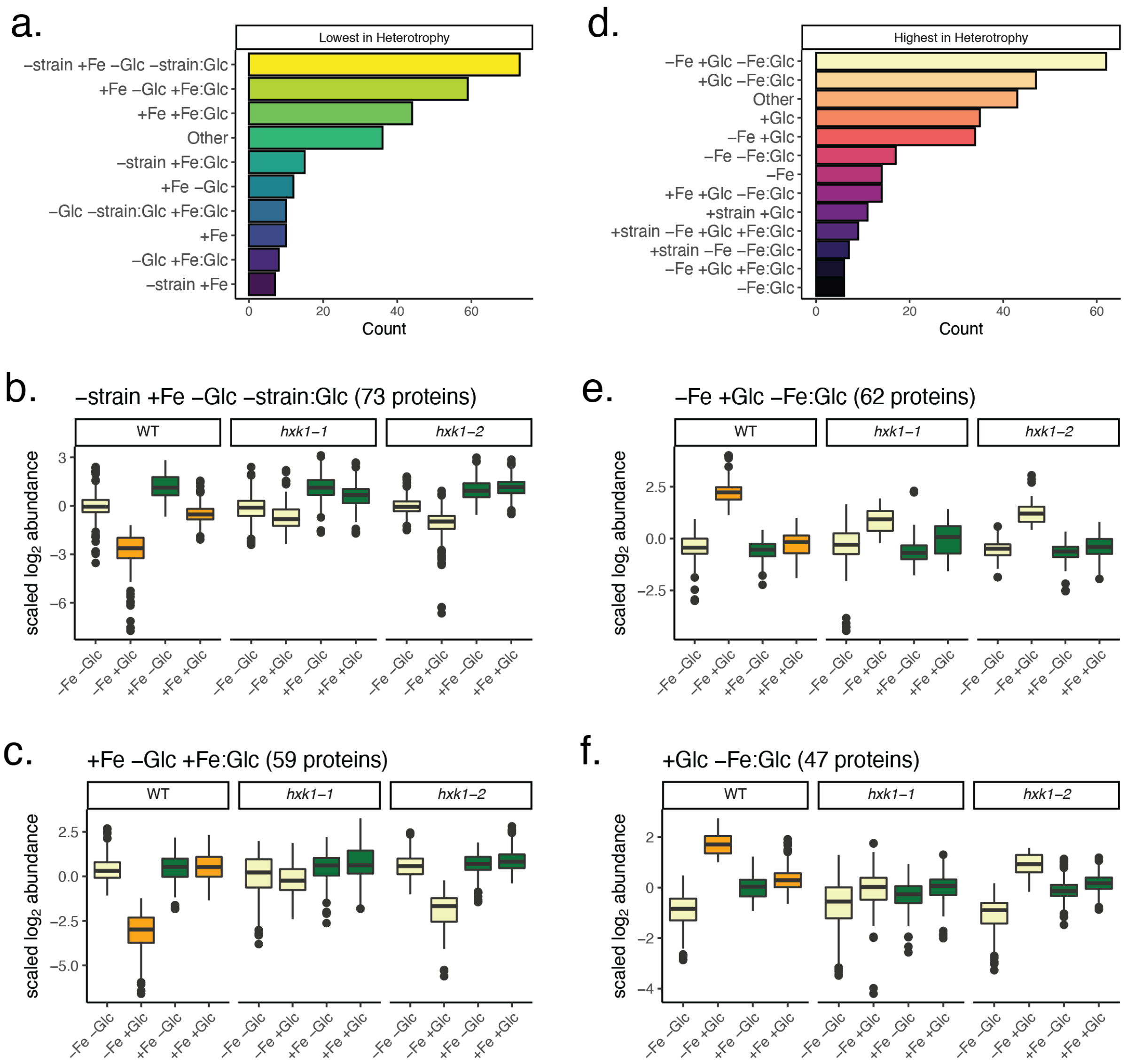


**Supplementary Fig. 8 Proteomics linear model categorization groups of unique to heterotrophy.**
The dominant equation subgroups assigned to the significant in heterotrophy proteins vary, showing proteins that are lowest-in-heterotrophy may have differing responsiveness to Fe, Glc, or strain. **a.** The lowest-in-heterotrophy proteins are most often categorized by subgroups "-strain +Fe −Glc +Fe:Glc" **(b)** or "+Fe −Glc +Fe:Glc" **(c)**. **d.** The highest-in-heterotrophy protein are most often categorized by subgroups "−Fe +Glc −Fe:Glc"**(e)** or "+Glc −Fe:Glc" **(f)**.

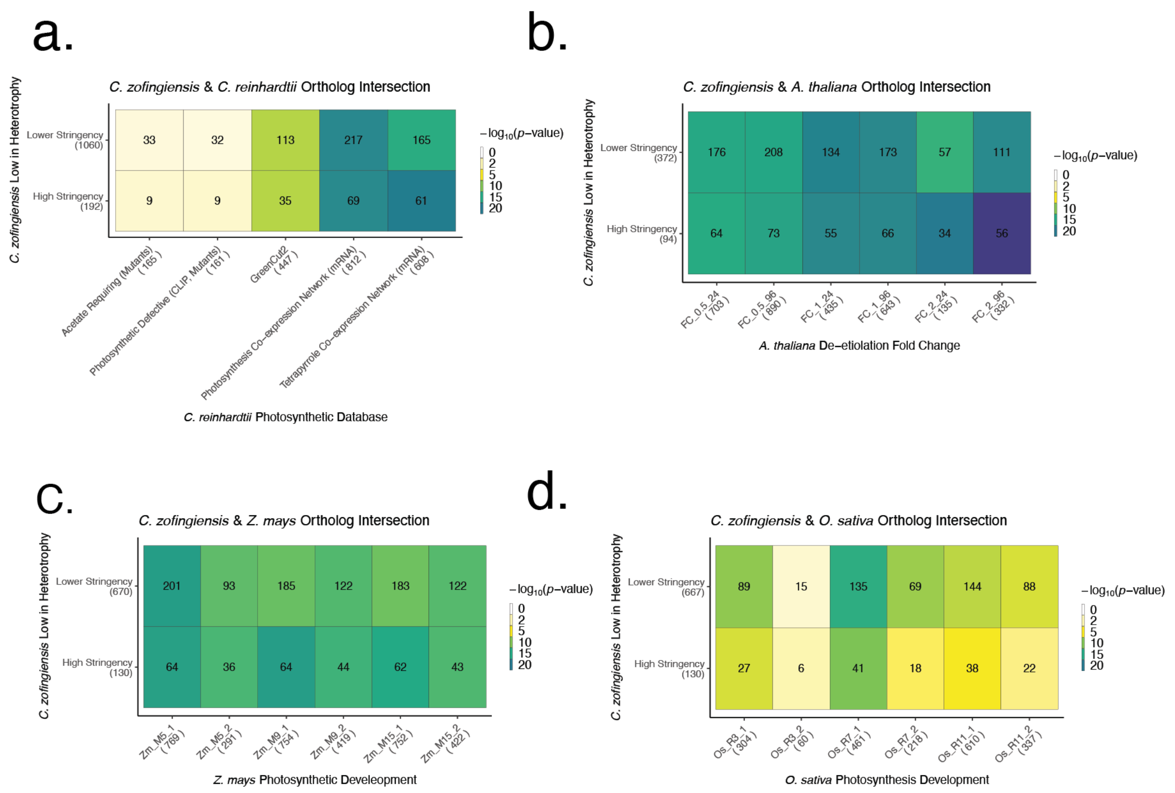


**Supplementary Fig. 9 Significance of ortholog overlap in photosynthetic regulation with reference organisms.**
**a.** Heatmap ortholog enrichment overlap analysis between the highly stringent and less stringent photosynthesis lists (y-axis) in *C. zofingiensis* and several photosynthesis associated lists developed in *C. reinhardtii*[8-11] (*x*-axis). The number of total distinct ortholog groups per photosynthesis gene list is in parenthesis on axes labels. The number of overlapping orthologs groups found in each species' gene list is written in heatmap block, which is colored by the p-value of overlap significance determined by the Fisher exact test. Only nucleus-encoded genes were included in analysis due to lack of *C. reinhardtii* plastid-encoded genes used in transcriptome and phylogenomic analysis. **b.** Significance heatmap of *C. zofingiensis* lowest in heterotrophy and their orthologs upregulated de-etiolation in the proteome measured in *A. thaliana*[12.] The x-axis considers various stringency of induction, with $log_2$ fold change (FC) greater than >0.5, >1, or >2 from the 0 h etiolated time point in both 24 and 96 h. Plastid encoded proteins are included in analysis. **c.** *C. zofingiensis* lowest in heterotrophy ortholog with *Zea mays* photosynthetic development upregulated along basipetal axis. M5, M9, and M15 refer to various photosynthetic tissues and their upregulation in expression compared to the undeveloped photosynthetic tissues below the ligule (M1)[13]. **d.** *C. zofingiensis* lowest in heterotrophy orthologs with *Oryza sativa* photosynthetic development along basipetal axis. Like in *Z. mays* R3, R7, and R11 are photosynthetically developed tissue whose upregulation in expression is compared to the undeveloped photosynthetic tissues below the ligule (R1)[13].

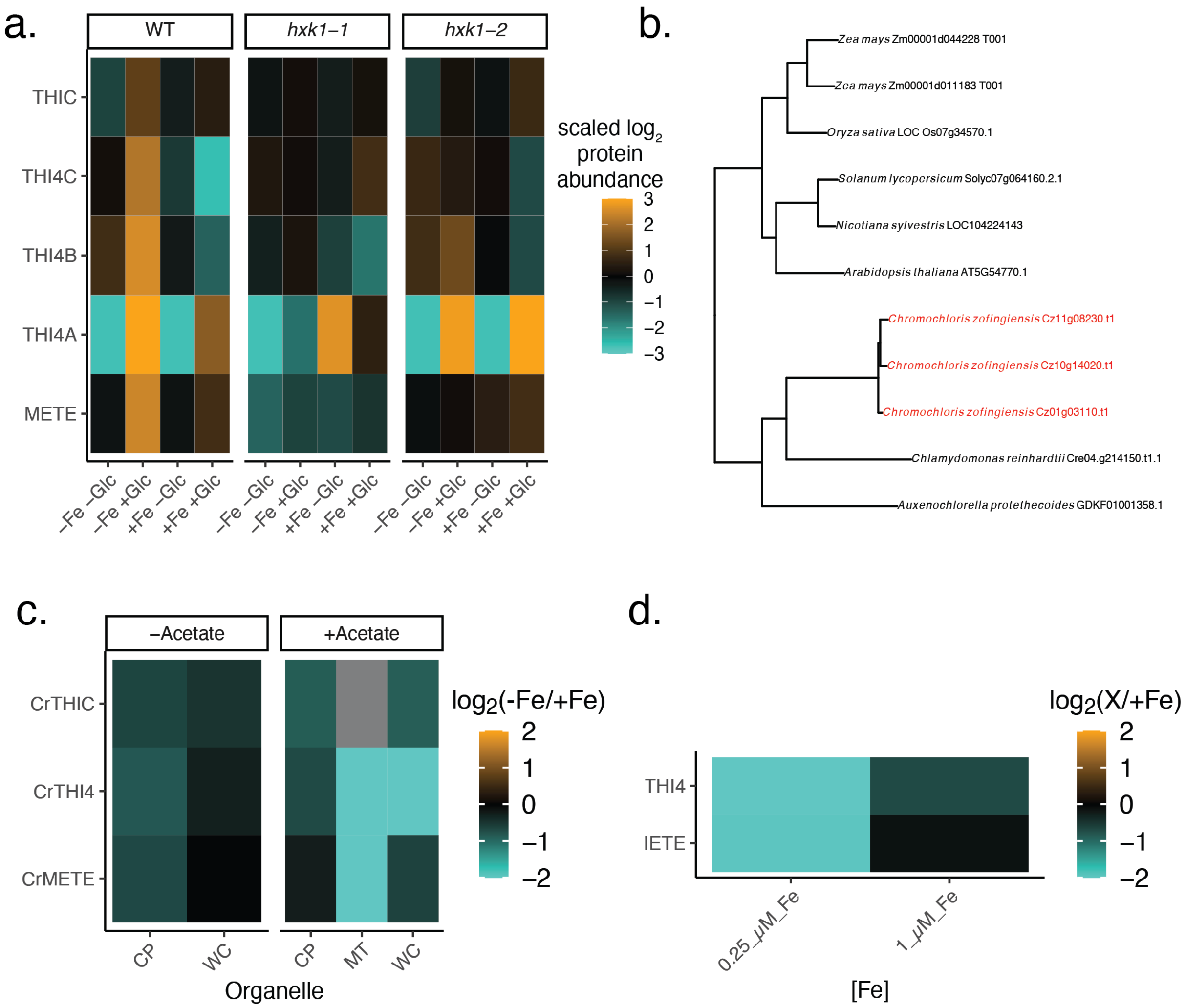


**Supplementary Fig. 10 Thiamine and methionine biosynthetic processes are upregulated in heterotrophy.**
**a.** Proteomics heatmap showing upregulation of THIC, all C. zofingiensis copies of THI4 (manually labeled A-C), and the cobalamin-independent methionine synthase (METE). **b.** Orthofinder2[14] produced gene tree of THI4 enzymes shows *C. zofingiensis* uniquely contains three copies of THI4 (labelled in red) **c.** Protein abundance of *C. reinhardtii* to −Fe for THIC, THI4, and METE in various organellar extracts (*x*-axis: chloroplast (CP), mitochondrion (MT) and whole cell (WC)) and +/- acetate. Data source: Supplemental Ref. 1. **d.** Soluble proteome extract of *C. reinhardtii* in 0.25 µM Fe (limited) and 1 µM Fe (deficient) $\log_2$ ratio over 20 µM Fe (replete). Data source: Supplemental Ref. 2 . METE and THI4 are highly depleted in 0.25 µM Fe. Cultures were grown in acetate and THIC abundance was not reported in publication.

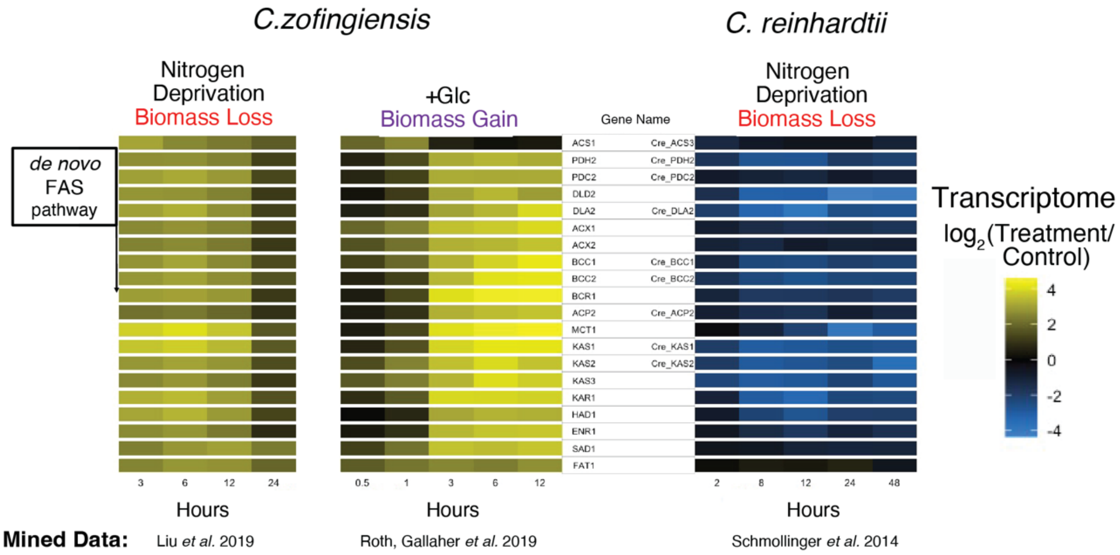


**Supplementary Fig. 11 Transcriptome time courses of the *de novo* fatty acid synthesis pathway in *C. zofingiensis* and *C. reinhardtii*.**

*C. zofingiensis* rapidly upregulates (yellow scale) a complete *dn*FAS pathway in both photoautotrophic nitrogen deprivation[15] and glucose addition[6], despite different impacts of these nutrients on overall biomass. Nitrogen deprivation in *C. reinhardtii*[16] leads to a concerted downregulation of this entire pathway (blue). Transcriptional data is $log_2$ normalized the ratio of time-controlled +Glc sample over condition not treated with Glc. Nitrogen conditions in both species are normalized to a 0 h timepoint.

#### Supplementary Table 1. AFS freeze-substitution procedure for cell suspensions.

| Step | Temp (ºC) | Time (h) | Gradient (ºC $h^{-1}$) |
|---|---|---|---|
| T1 | -90 | 48 | |
| S1 | | 6 | 5 |
| T2 | -60 | 20 | |
| S2 | | 16 | 5 |
| T3 | 20 | 24 | |

Substitution medium at T1-T3 was 1% glutaraldehyde and 0.1% tannic acid in anhydrous acetone.
