## Supplementary figures and images for "Iron rescues glucose-mediated photosynthesis repression during lipid accumulation in the green alga *Chromochloris zofingiensis*"

### 20210429_D9_27-3_+Fe-Glc.jpg

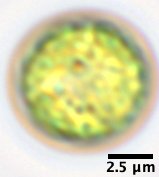

### 20210429_D918-1_-Fe-Glc.jpg

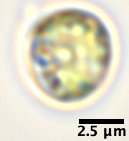

### 20210429_D919-1_-Fe-Glc.jpg

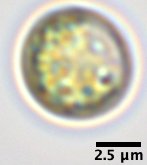

### 20210429_D921-3_-Fe+Glc.jpg

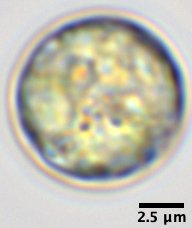

### 20210429_D927-3_+Fe-Glc.jpg

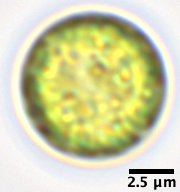

### 20210429_D931-3_+Fe+Glc.jpg

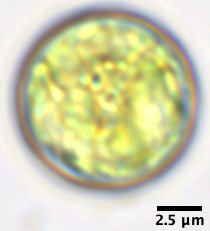

### 20210429_WT01_-Fe-Glc.jpg

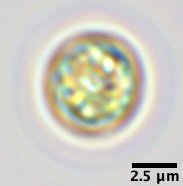

### 20210429_WT05_-Fe+Glc.jpg

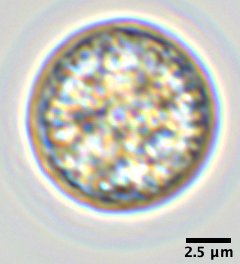

### 20210429_WT09_+Fe-Glc.jpg

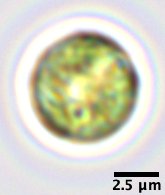

### 20210429_WT15_+Fe+Glc.jpg

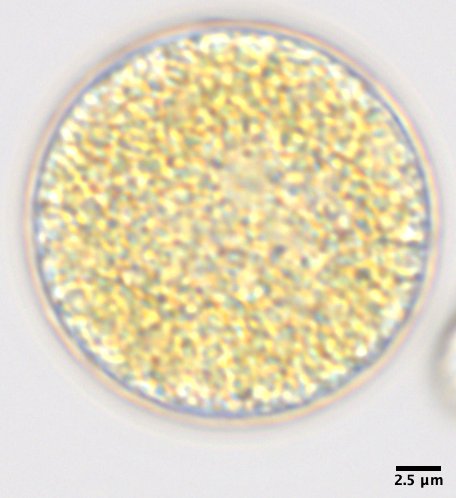

### 20210501_D633_-Fe-Glc.jpg

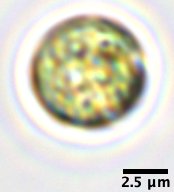

### 20210501_D638_-Fe+Glc.jpg

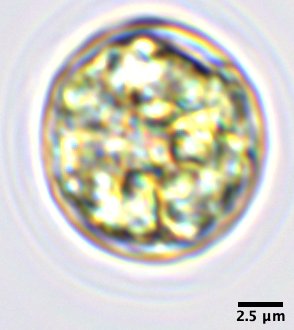

### 20210501_D641_+Fe-Glc.jpg

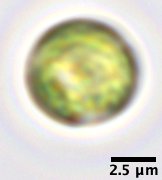

### 20210501_D645-3_+Fe+Glc.jpg

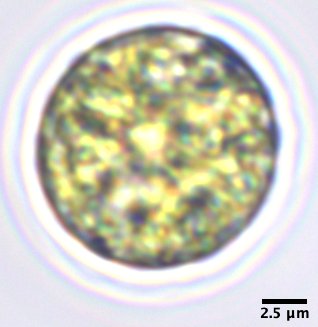

### A-wt17_s-17.jpg

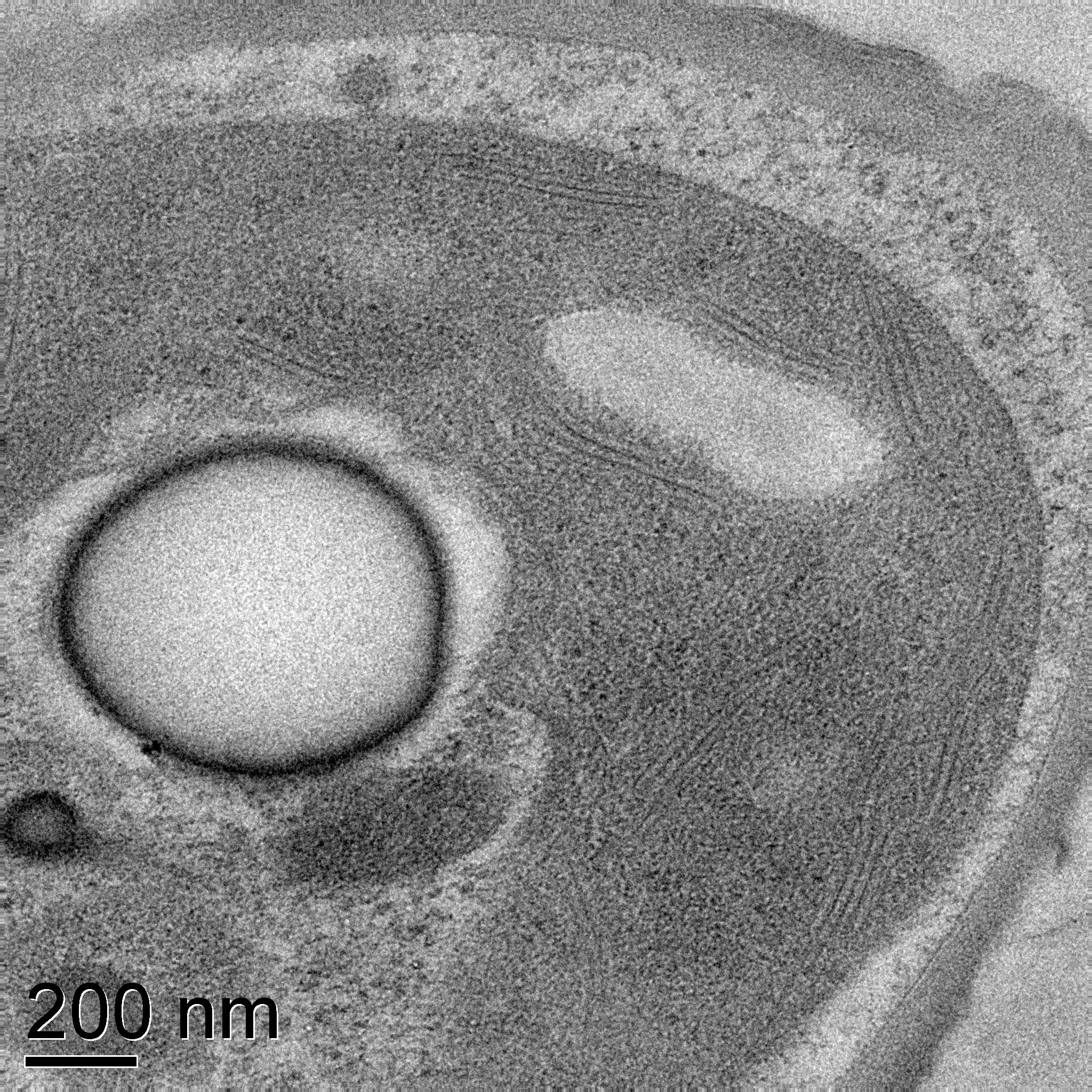

### A-wt17_s-23.jpg

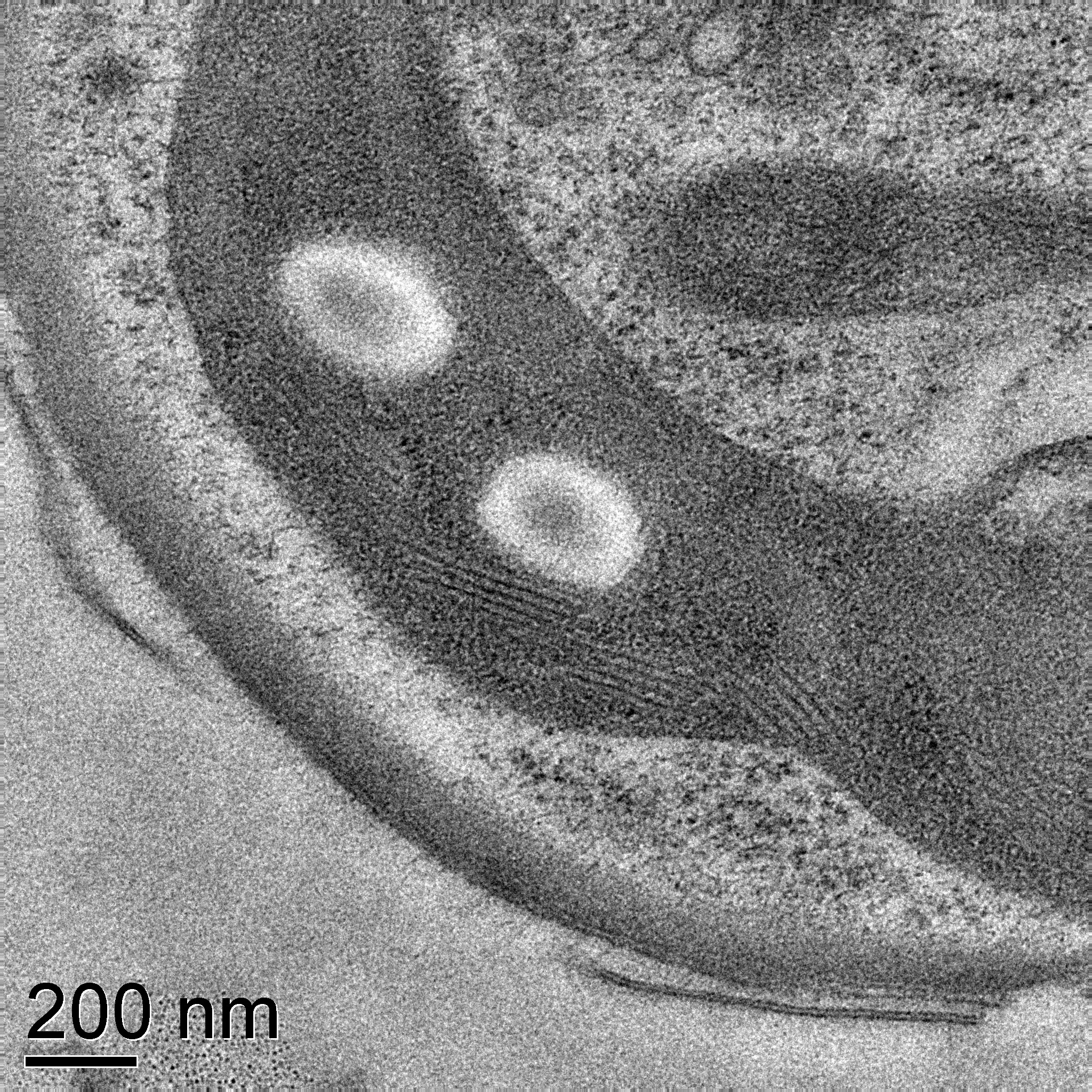

### A-wt17_s-34.jpg

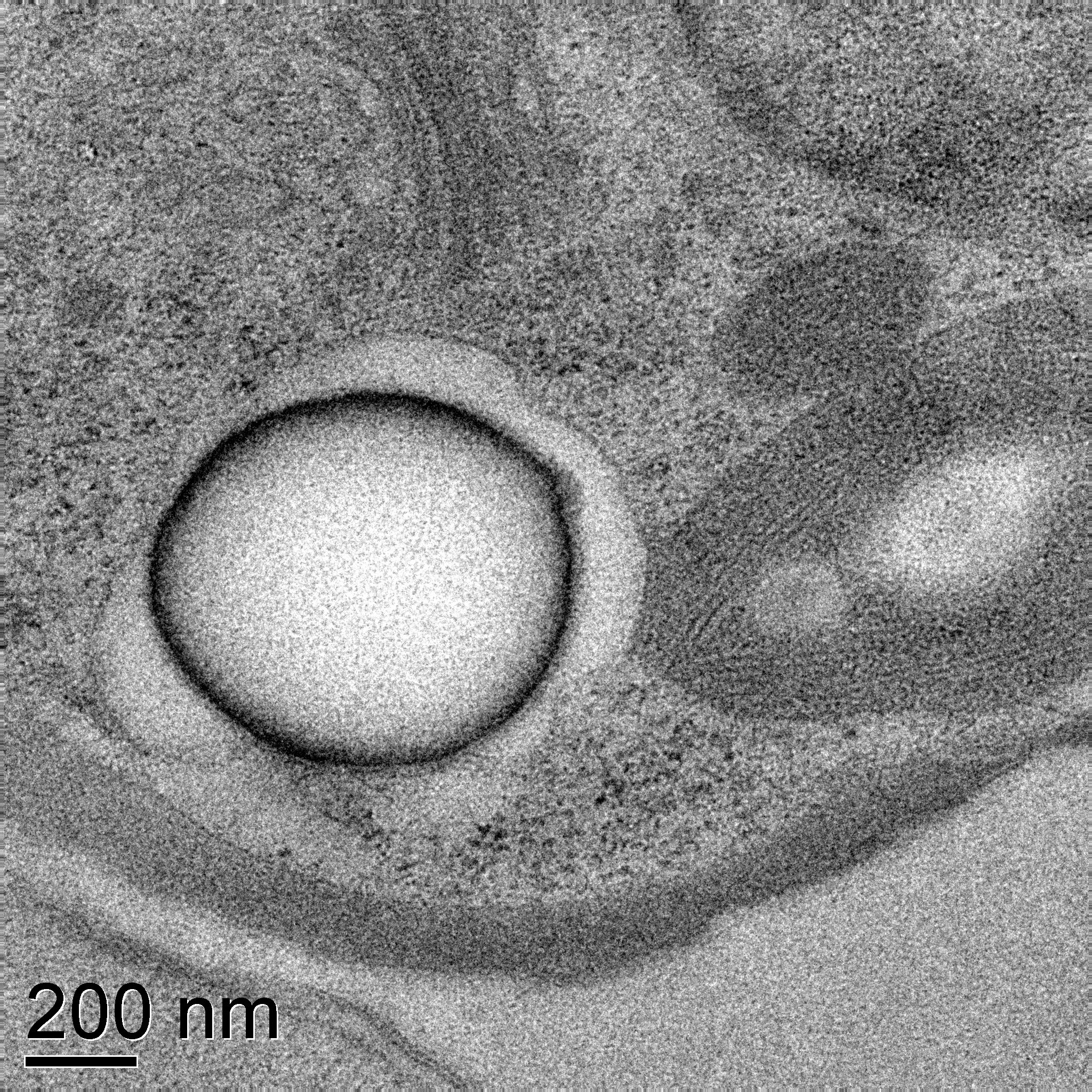

### B-wt25s-5.jpg

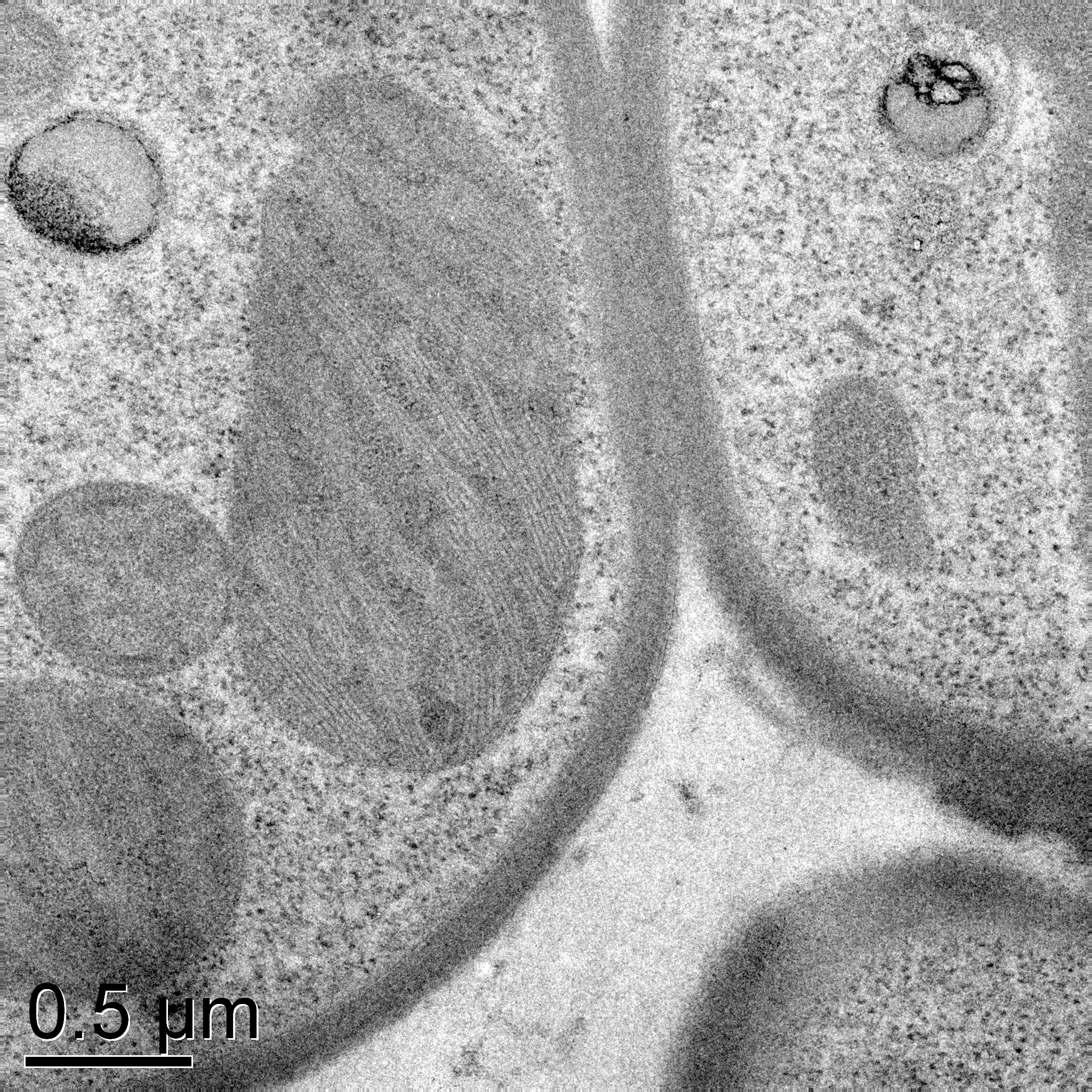

### B-wt25s-12.jpg

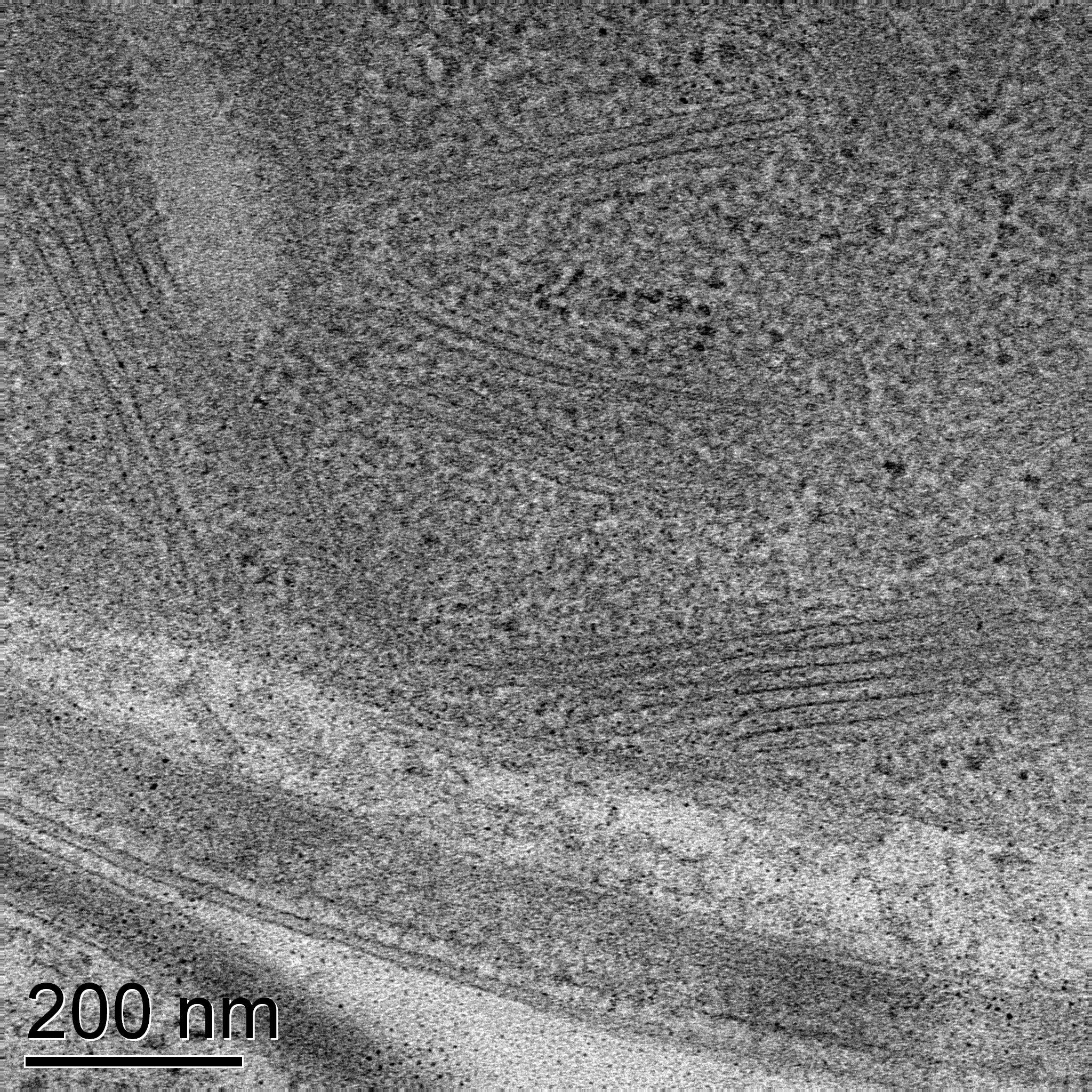

### B-wt25s-26.jpg

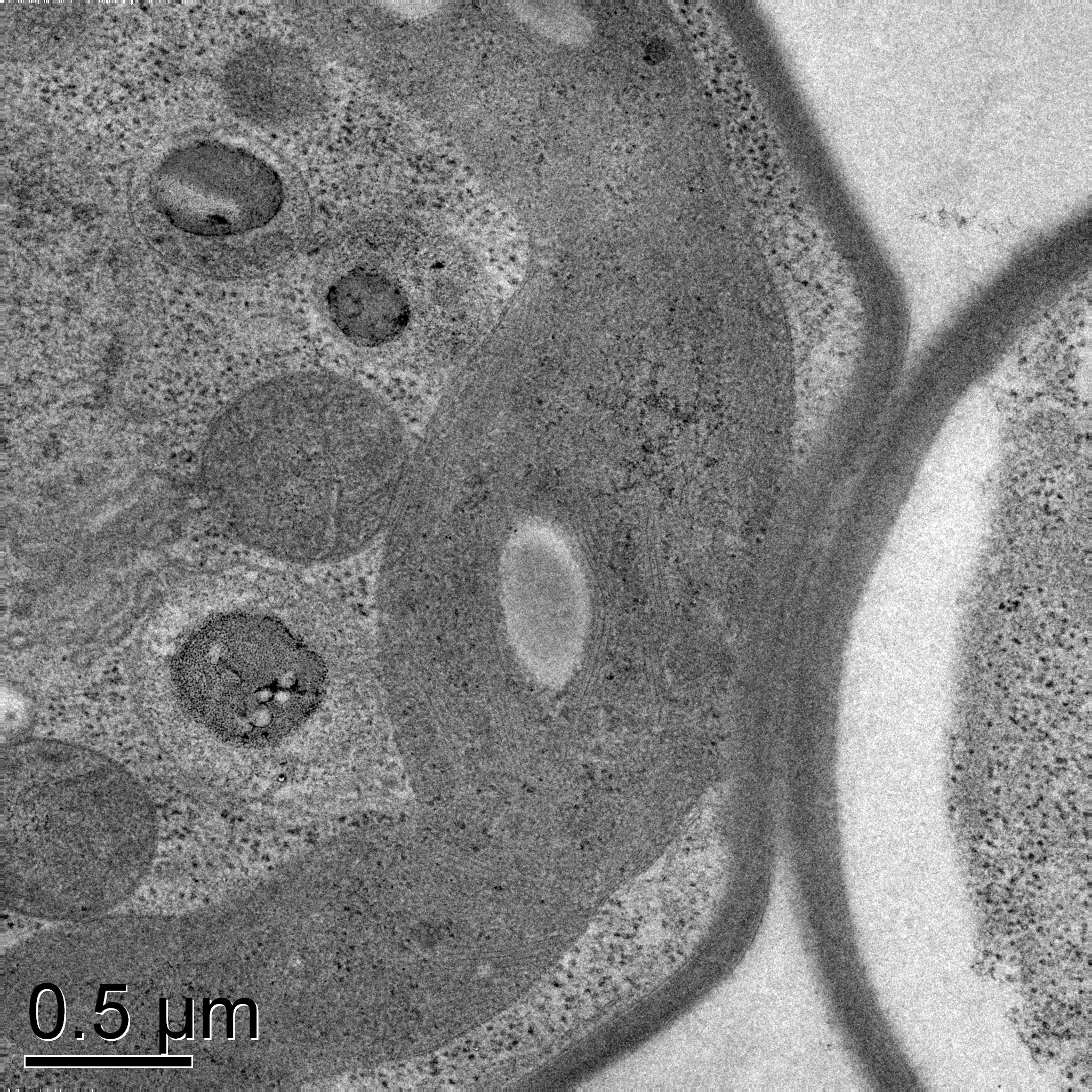

### C-wt21_-Fe-5.jpg

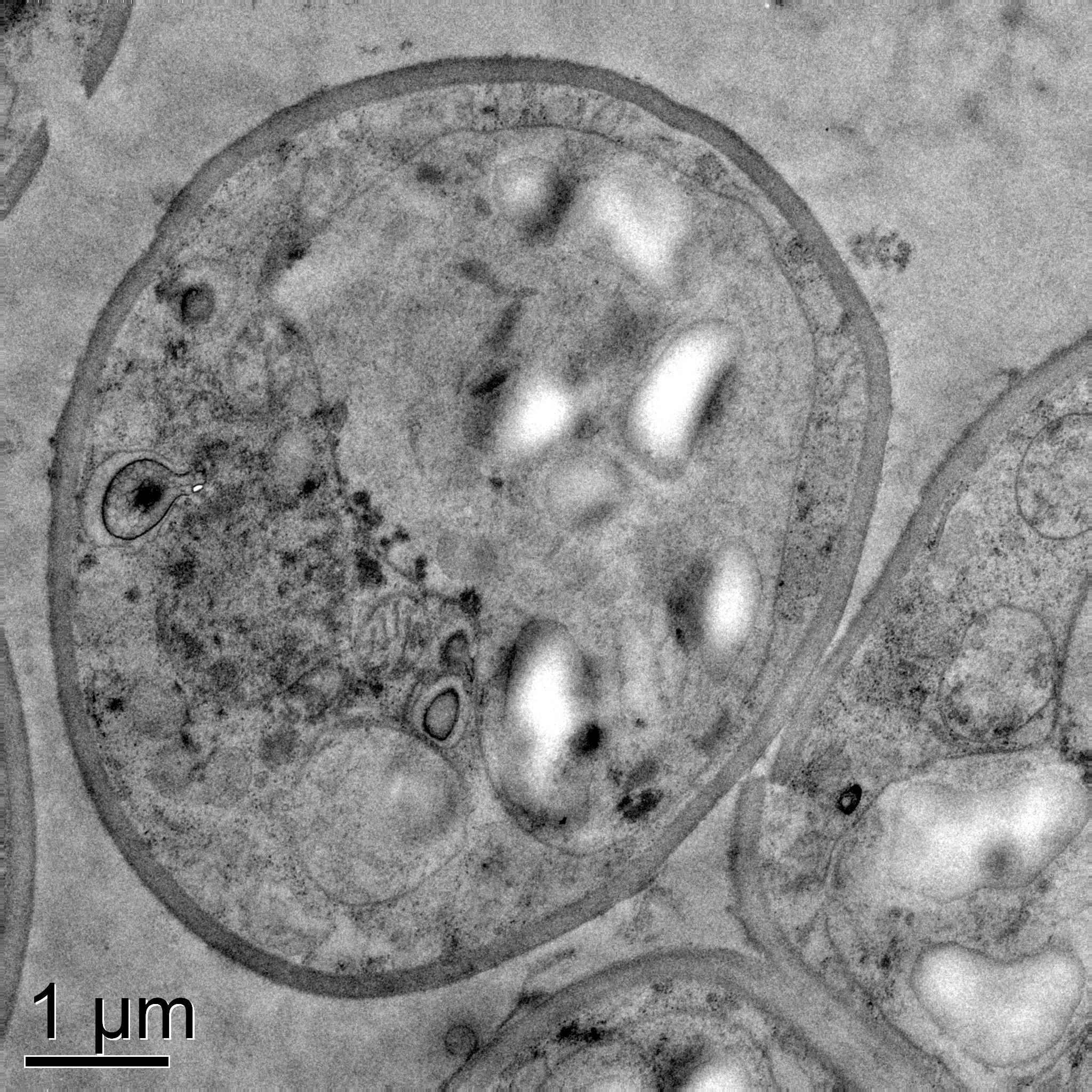

### C-wt21_-Fe-33.jpg

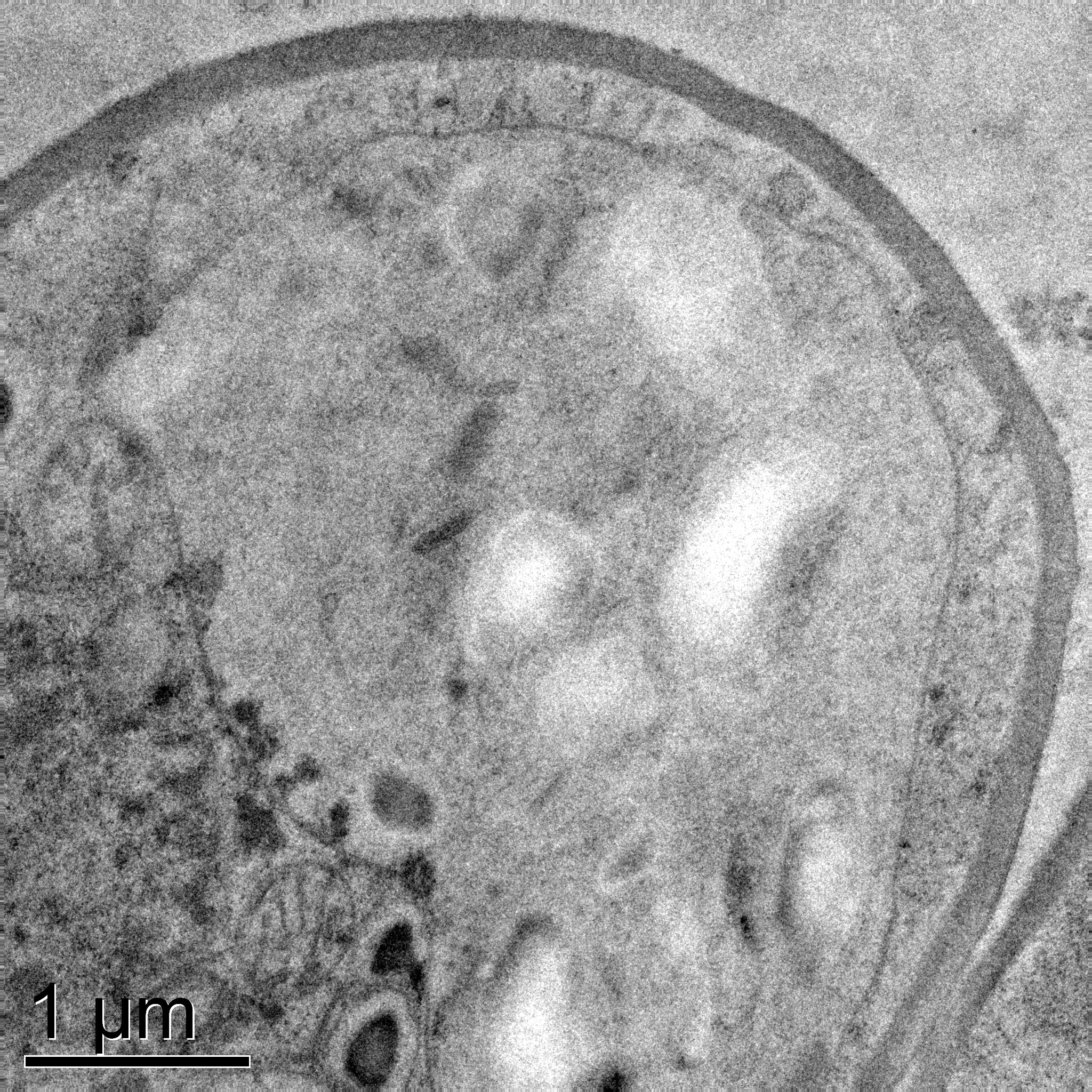

### C-wt21_-Fe-50.jpg

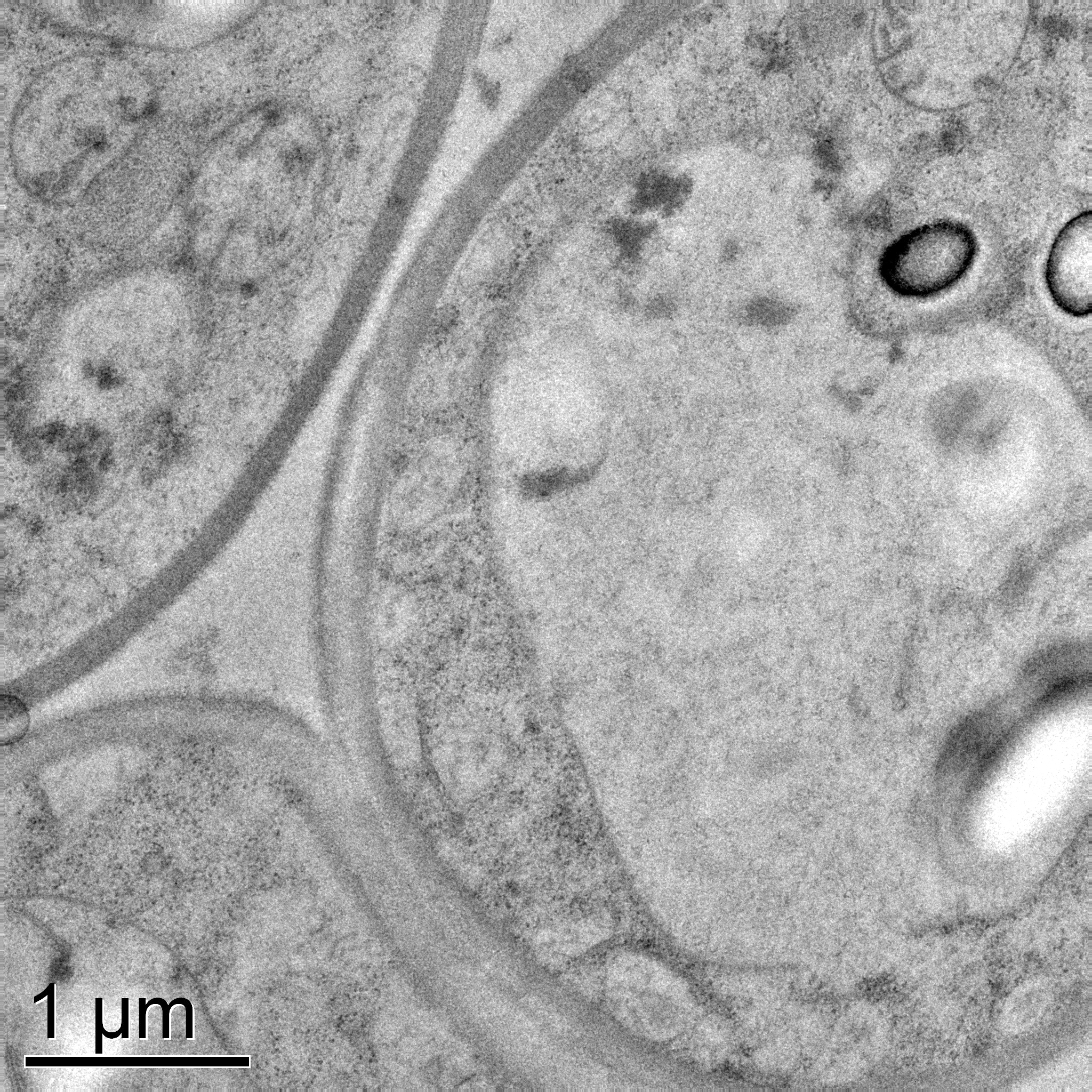

### D-wt29_+Gl+Fe-7.jpg

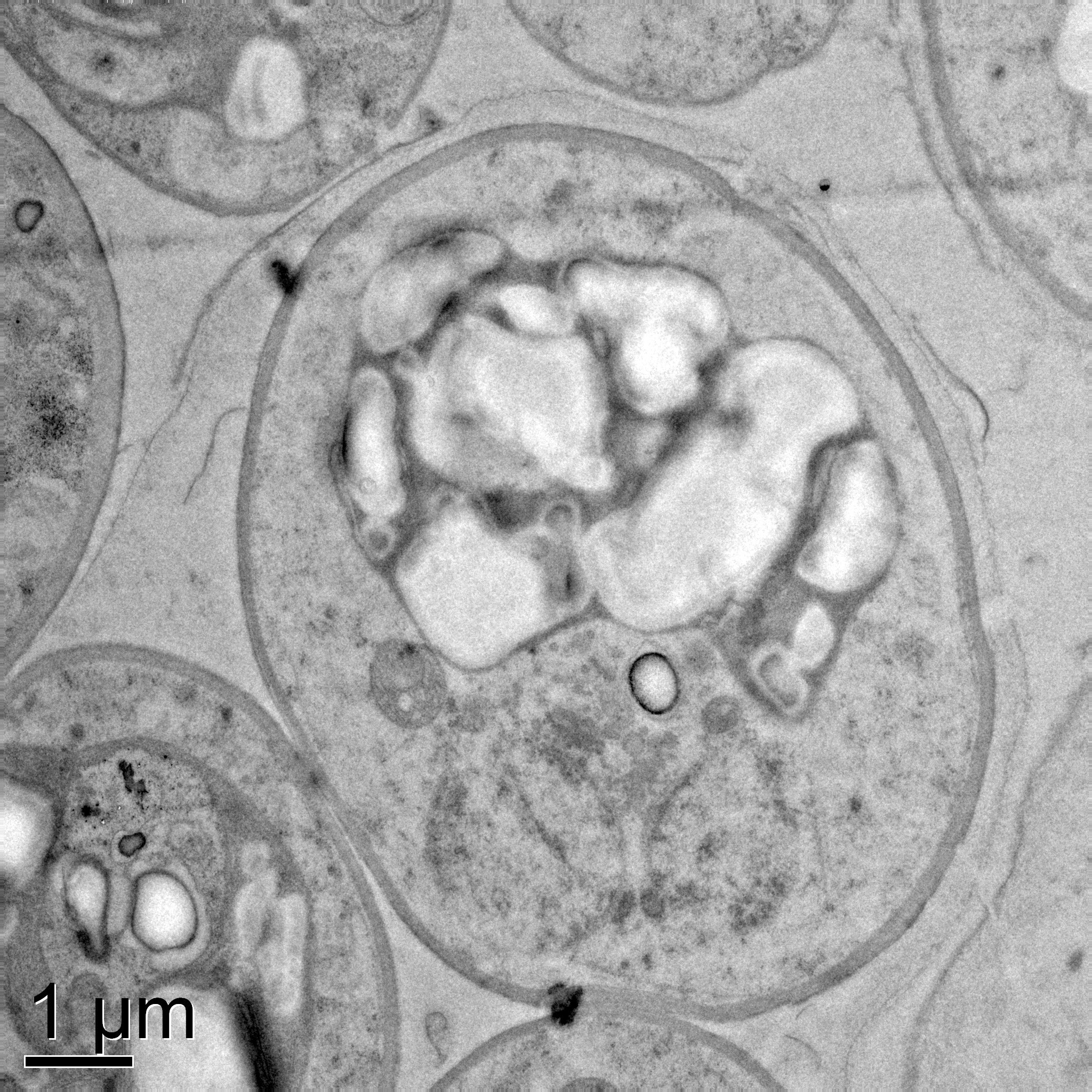

### D-wt29_+Gl+Fe-17.jpg

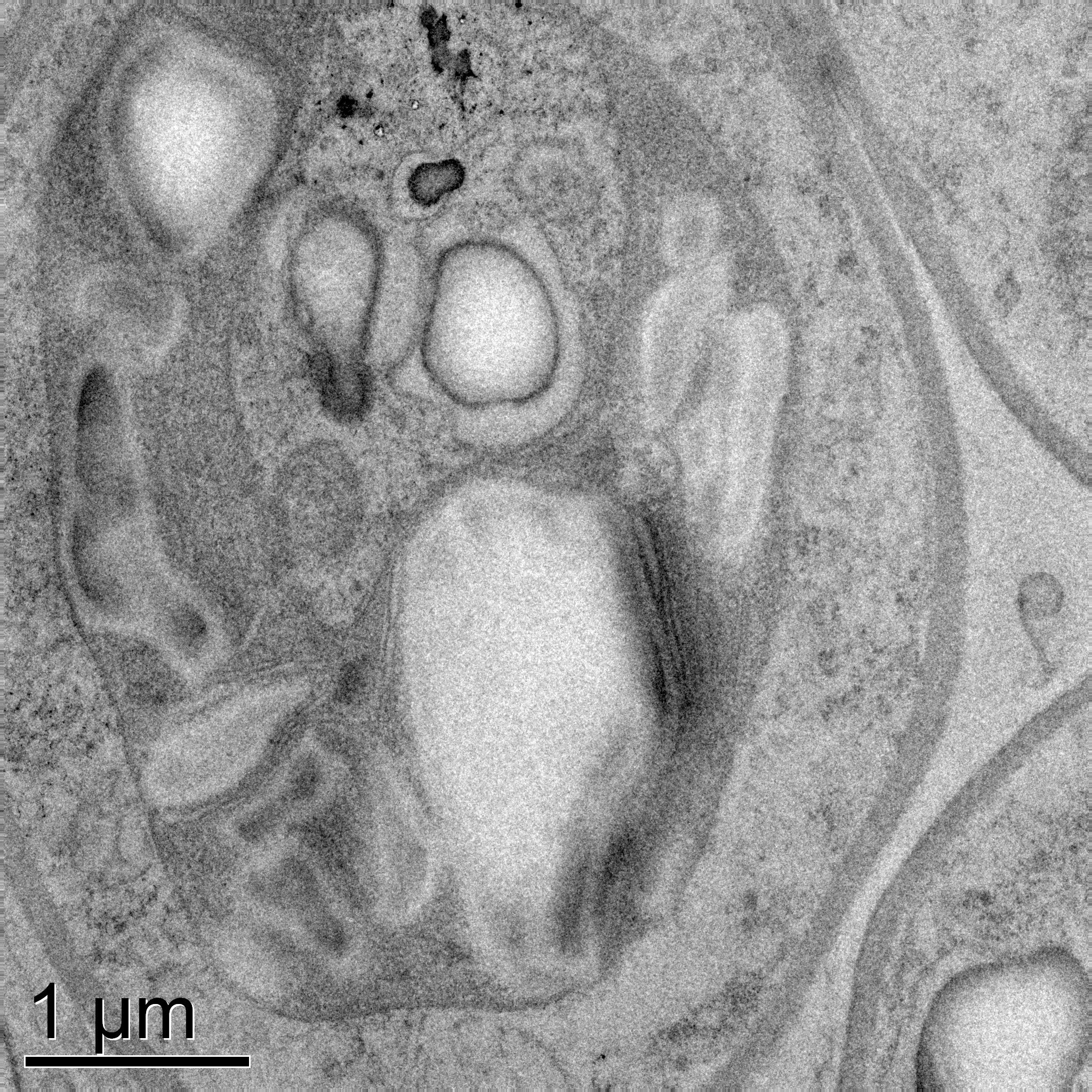

### D-wt29_+Gl+Fe-21.jpg

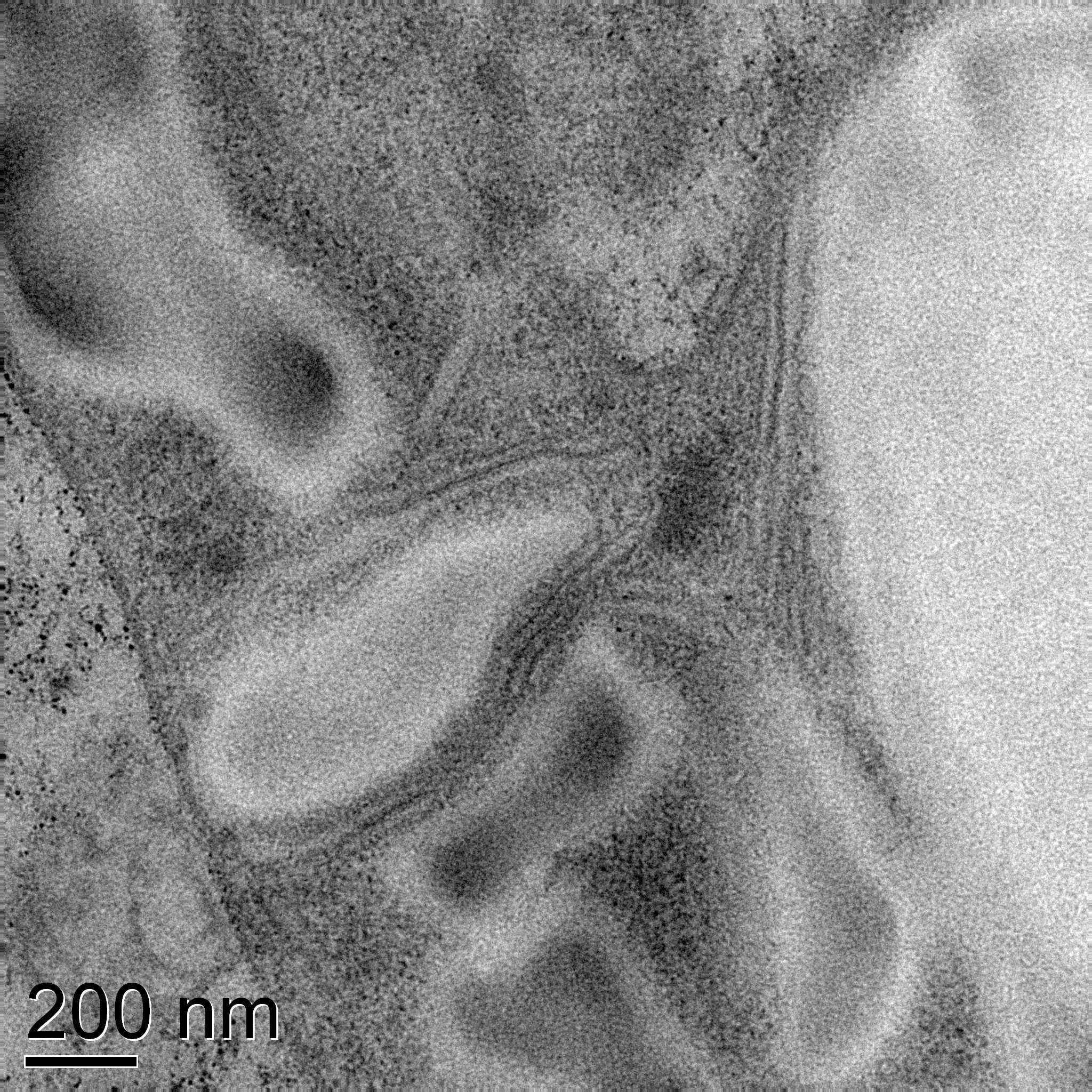

### Fig.2e.Uncropped_TLC_UV_withreplicates Annotated.jpeg

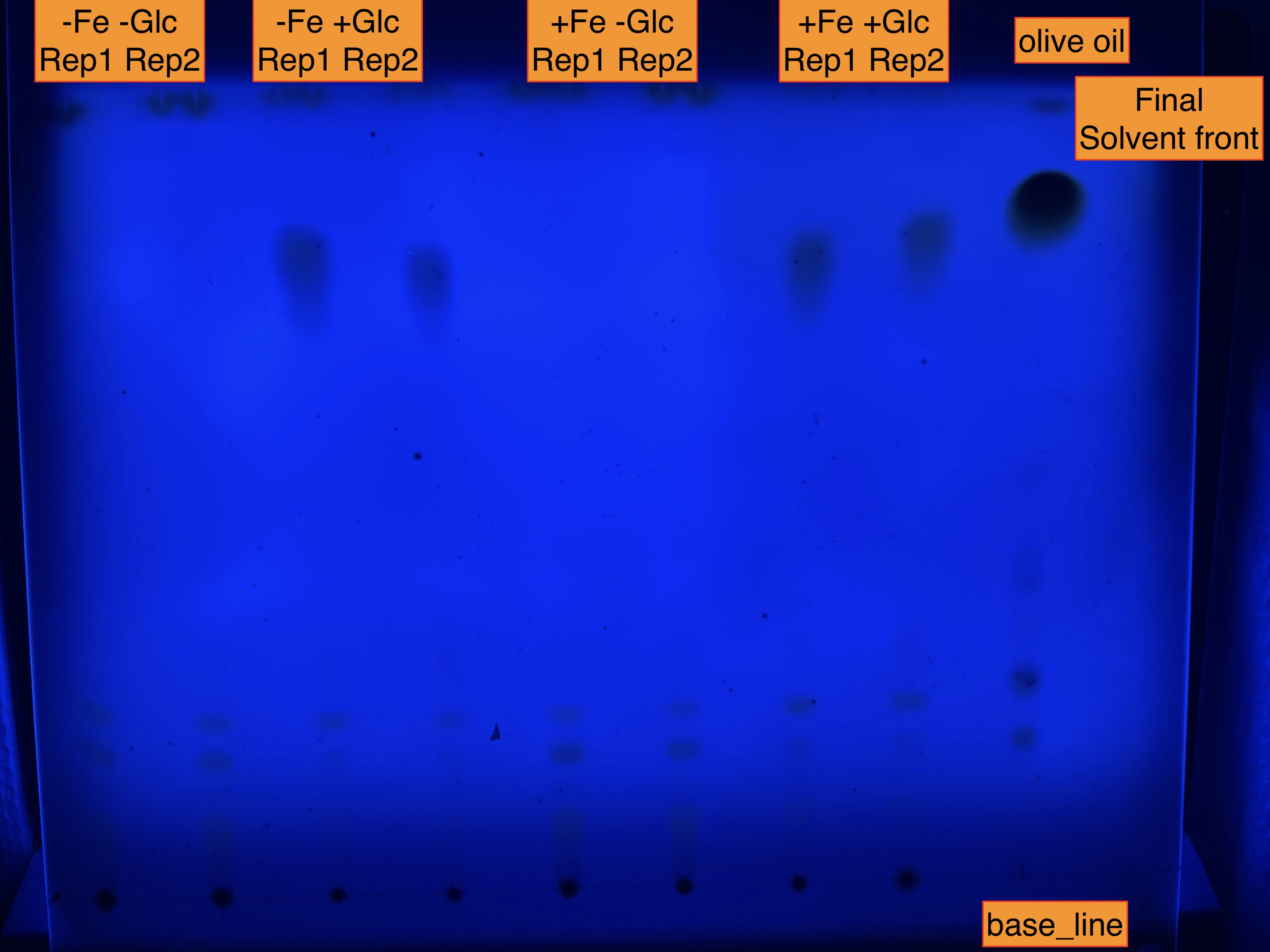

### Fig.2e.Uncropped_TLC_UV_withreplicates.jpg

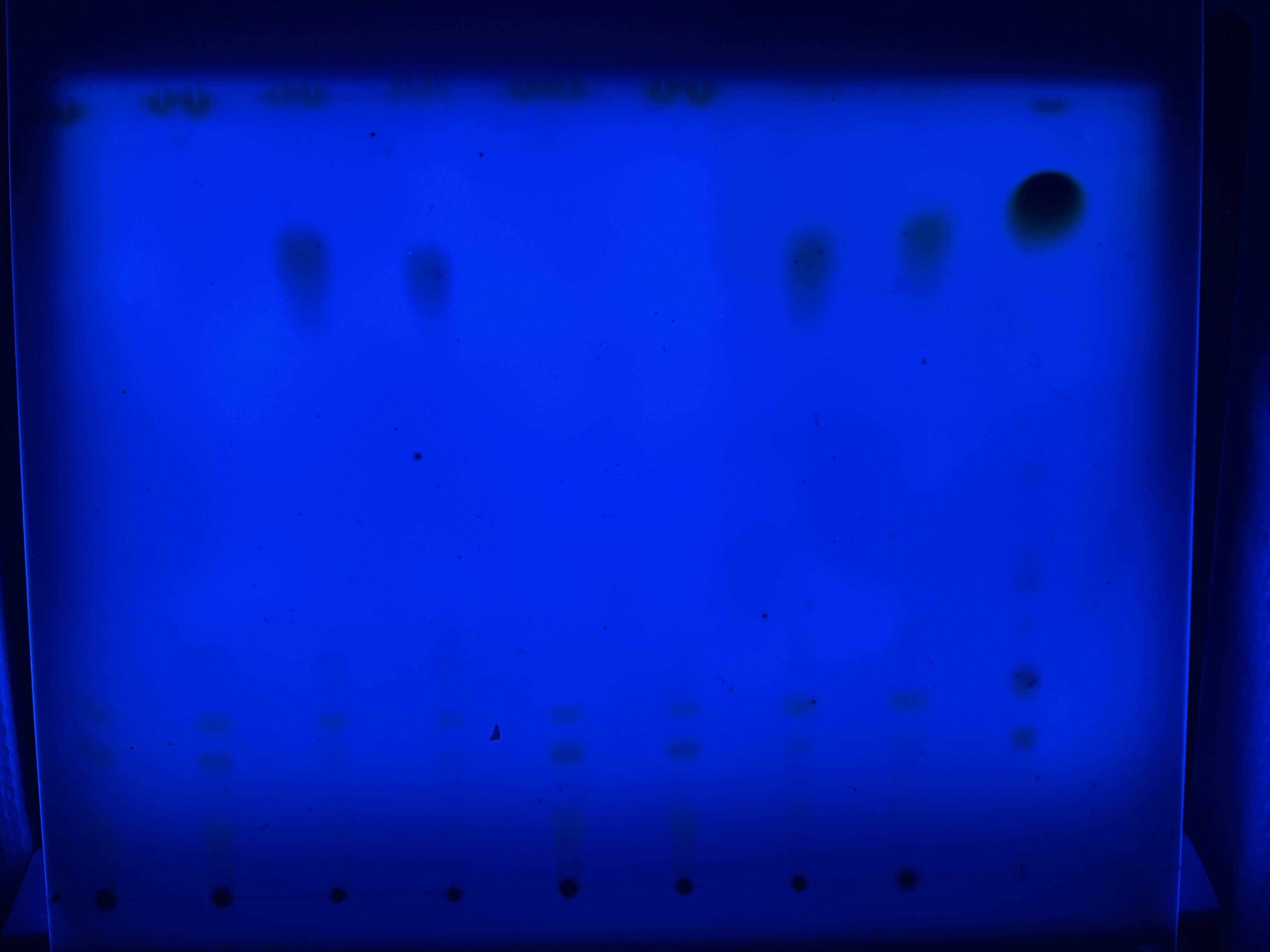

### Fig.7a_Uncropped_TLC_UV.jpg

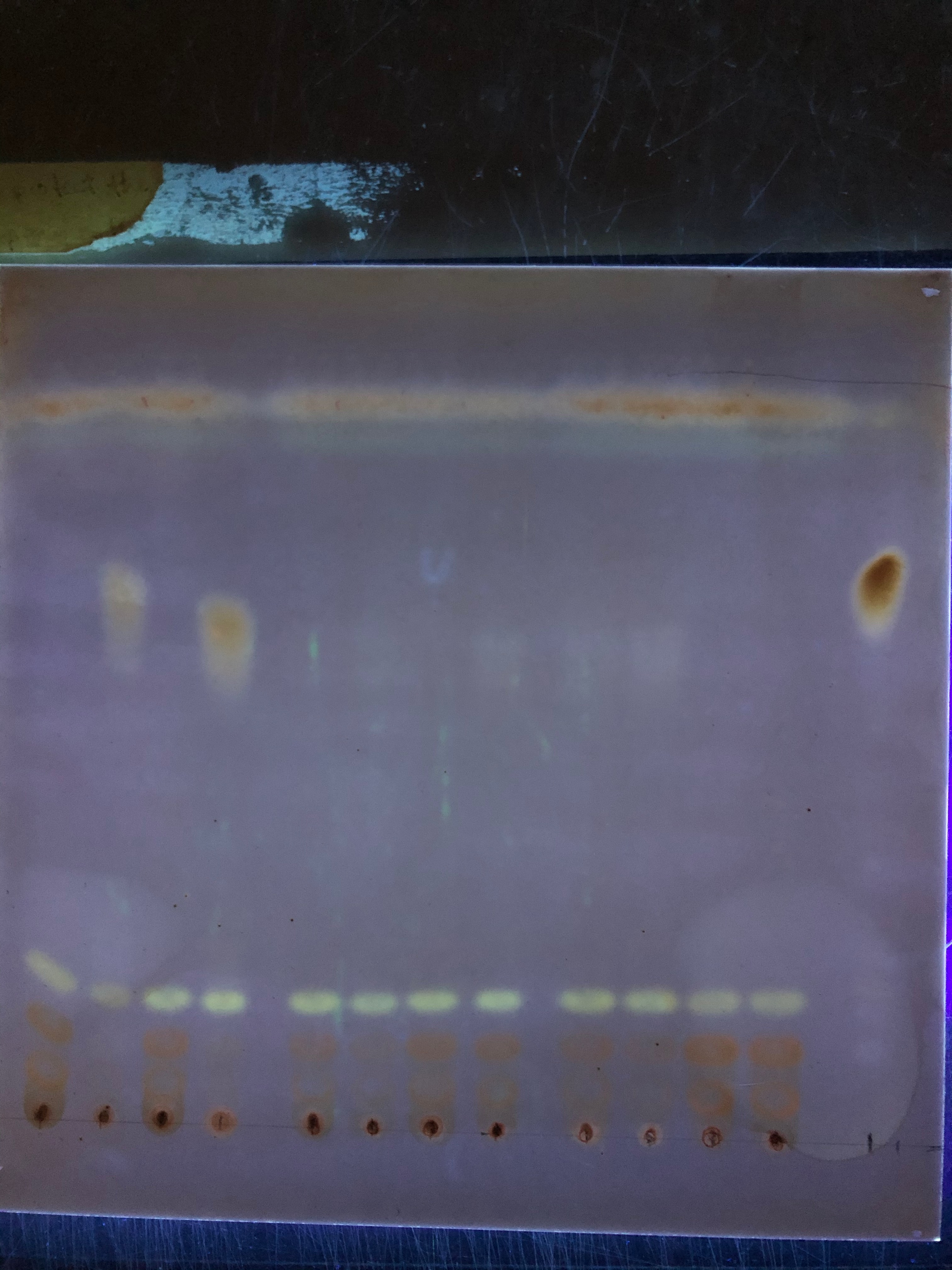
